# Optimized parameters for Cas9 CRISPR interference library design

**DOI:** 10.64898/2026.02.25.708018

**Authors:** Smriti Srikanth, Fengyi Zheng, Laura M Drepanos, Spencer T Shah, Eleanor G Kaplan, Daytan Gibson, Grace O Lynch, Allison T Uebele, Ganna Reint, Sarra Merzouk, John G Doench

## Abstract

CRISPR interference (CRISPRi) is a powerful technology for studying loss-of-function phenotypes, enabling transient and reversible control of gene expression without the introduction of double-stranded DNA breaks. The cost of conducting large-scale CRISPR screens necessitates the selection of effective and specific sgRNAs for the design of compact libraries. While several genome-wide Cas9 CRISPRi libraries have been created, updates to transcript annotations, the generation of higher-resolution chromatin accessibility datasets and the development of newer on-target prediction models motivate an updated CRISPRi library design approach. Here, we generate large CRISPRi datasets tiling essential and nonessential genes. We compare the performance of multiple KRAB domain systems, develop an updated CRISPRi-specific on-target scoring scheme, and quantitatively characterize off-target effects associated with seed sequence patterns. We leverage these findings to design an optimized Cas9 CRISPRi library, Katsano, and validate its performance with genome-wide viability screens.

## INTRODUCTION

Pooled CRISPR screening platforms offer a powerful approach to investigating gene function at scale.(1, 2) CRISPR interference (CRISPRi), which typically employs a nuclease-inactive *S. pyogenes* Cas9 (dCas9) tethered to a repressive domain, presents a useful alternative to CRISPR knockout (CRISPRko) for performing loss-of-function genetic screens.(3–7) Unlike CRISPRko, CRISPRi offers the advantage of not introducing double-stranded DNA breaks, thereby reducing associated toxicity unrelated to target gene function. Furthermore, its transient nature enables titration of gene expression to study gene dosage phenotypes.(8)

The high cost of large-scale screens necessitates the design of compact CRISPR libraries, particularly with recent advances in high-dimensional screening readouts.(9–13) When employing fewer guides per gene, it is increasingly important to select those that effectively perturb the target; this has motivated efforts toward accurate prediction of sgRNA activity.(14) In addition to intrinsic sequence properties, robust transcriptional repression with CRISPRi relies upon guide RNA location relative to the target transcription start site (TSS), as well as sustained chromatin accessibility.(5, 6, 15, 16) With these considerations in mind, several genome-wide Cas9 CRISPRi libraries have been curated to facilitate functional interrogation of the human genome. Human CRISPRi v1 (hCRISPRiv1) includes 10 sgRNAs per protein-coding gene, selected from a TSS-proximal window using simple sequence-based rules for off-target avoidance.(5) Human CRISPRi v2 (hCRISPRiv2) built upon this effort, integrating TSS proximity, sequence characteristics, and chromatin accessibility features into a machine learning model to select 10 sgRNAs per target with high predicted on-target efficacy.(6) Importantly, this update also featured the use of Cap Analysis of Gene Expression (CAGE) data generated by the FANTOM consortium for improved TSS annotation.(15, 17) We prioritized similar criteria for the design of Dolcetto, which includes 6 sgRNAs per gene.(7)

Since the creation of these libraries, the field of functional genomics has seen several advances. Updates to genome annotations reduce the coverage of existing libraries over time, and improved TSS annotations enable more informed guide selection.(14) In particular, the curation of the Matched Annotation from NCBI and EMBL-EBI (MANE) Select set of transcripts now provides a standardized, high-confidence human reference transcriptome.(18) The Encyclopedia of DNA Elements (ENCODE) consortium has continued to generate and expand access to high-resolution chromatin accessibility datasets.(19) Additionally, for the selection of guides in the CRISPRko context, we recently developed Rule Set 3 Sequence, an optimized sequence-based on-target prediction model that has yet to be applied to genome-wide CRISPRi library design.(20) Improvements to CRISPRi technology itself, including the characterization of new Krüppel-associated box (KRAB) domains and vector architectures, further encouraged re-optimization of our CRISPRi systems.(21–23)

We executed large scale CRISPRi screens tiling the promoter regions of essential and nonessential genes, with which we establish comparable performance of multiple repressive domains and demonstrate that dCas9 N-terminal KRAB fusions are generally more effective than C-terminal fusions. Leveraging these data in combination with previously published tiling datasets, we elucidate features associated with guide efficacy and integrate them into an updated on-target model, Rule Set 3 interference (RS3i). We additionally identify sgRNA seed sequence characteristics that promote off-target behavior that differ substantially from the off-target profile of CRISPR knockout guides and corroborate seed-driven effects observed in prior studies.(24–27) We apply these insights to the design of a new genome-wide Cas9 CRISPRi library, Katsano, which demonstrates substantially improved screening performance as compared to previous libraries.

## RESULTS

### Design and evaluation of large-scale CRISPRi tiling screens

To investigate factors influencing CRISPRi sgRNA on-target and off-target activity, we designed a library targeting 201 essential and 198 non-essential genes previously targeted in our Rule Set 3 Validation Cas9 CRISPRko tiling screens.(20) The library includes guides tiling 1,000 base pairs upstream and downstream of the MANE Select TSS of each gene, comprising a total of 108,574 sgRNAs (Figure 1a).

**Figure 1.**
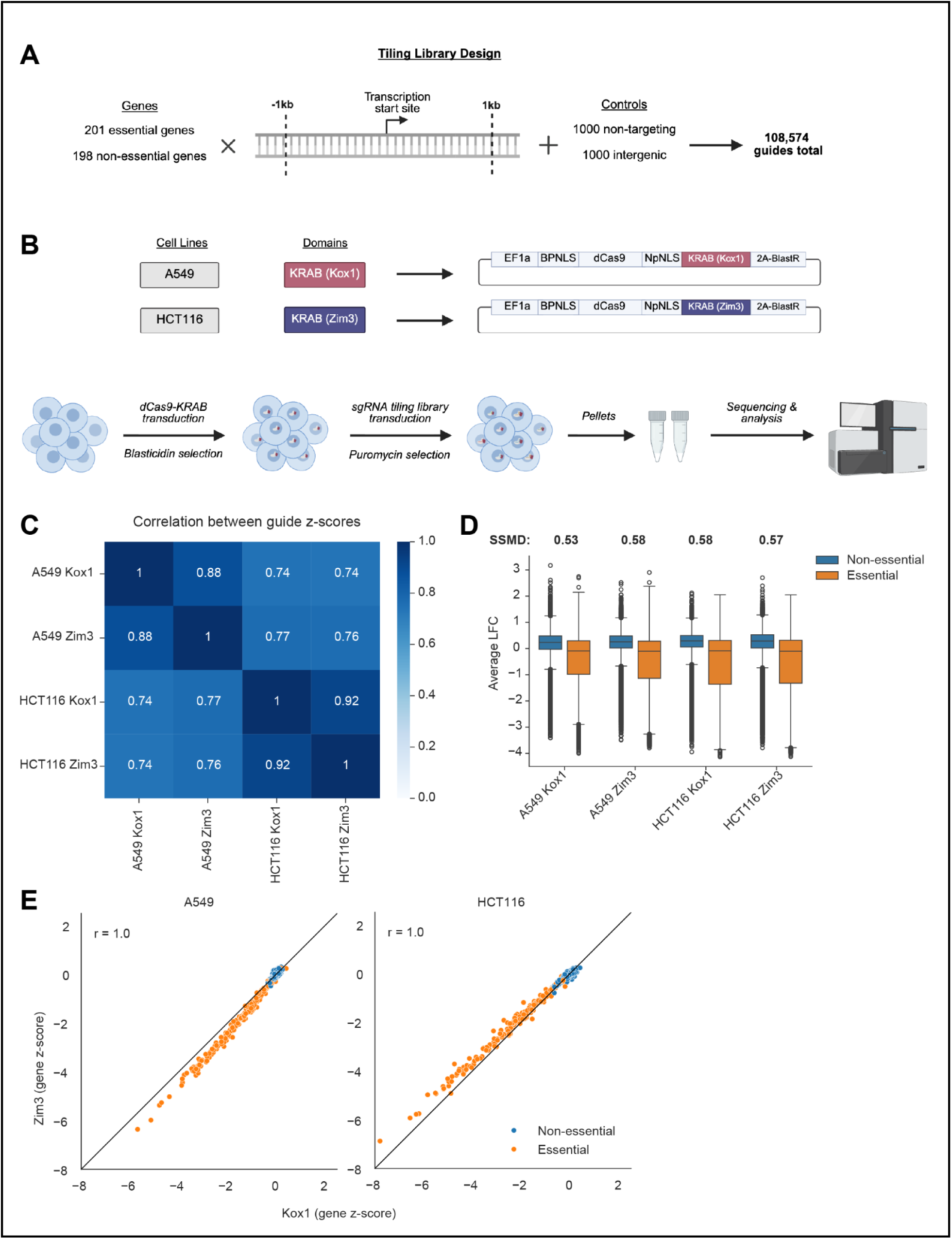
Design and evaluation of large-scale CRISPRi tiling screens. (A) Schematic illustrating design and composition of tiling library. (B) Schematic illustrating vector architecture and screening approach of tiling library. (C) Heatmap of Pearson correlation coefficients between guide-level z-scores in each screen performed. Z-scores were computed by averaging log-fold changes across replicates and z-scoring relative to intergenic controls. (D) Boxplot showing average log-fold changes of guides in screens performed in each cell line with each KRAB domain. Gene essentiality is indicated by color. Strictly standardized mean difference (SSMD) between non-essential and essential targeting LFCs in each screen is annotated above corresponding boxplots. Boxes show 25th (Q1), 50th (median), and 75th (Q3) percentiles, while whiskers show Q1 - 1.5*IQR and Q3 + 1.5*IQR (where IQR is Q3 - Q1). (E) Scatterplots of gene-level z-scores in screens performed with the Kox1 KRAB domain (x-axes) and the Zim3 KRAB domain (y-axes) in A549 (left) and HCT116 cell lines. Gene essentiality is indicated by color. Line of equality is shown as a solid black line. Gene-level z-scores were computed by averaging guide-level z-scores of guides targeting the same gene.

Prior studies have suggested that the KRAB domain from ZIM3 enables more potent transcriptional repression than that from KOX1 (hereafter abbreviated as Zim3 and Kox1 respectively).(21, 23) The two domains share 61% amino acid sequence identity, suggesting a moderate level of structural and functional similarity. We leveraged our tiling library to investigate the relative performance of Kox1 and Zim3 at scale. Using Fragmid(28), our modular vector assembly webtool, we generated separate constructs with each of the domains tethered to the C-terminus of dCas9 and screened the guide library in A549 and HCT116 cell lines expressing each construct (Figure 1b). These cell lines have been extensively characterized by the ENCODE consortium, facilitating downstream analysis of epigenetic guide efficacy determinants.(19)

The screens were performed in replicates and we observed strong reproducibility (Pearson R = 0.96-0.97, Supplementary Figure 1a). The results were also highly consistent across both KRAB domains and cell lines, with similar separation between nonessential and essential targeting sgRNAs in all four screens (Figure 1c-d). Interestingly, Zim3 performed slightly better in A549s while the Kox1 domain was marginally more effective in HCT116s, as indicated by more negative essential gene z-scores (Figure 1e). Together, these results demonstrate comparable performance of Kox1 and Zim3 in multiple cell lines.

### Design and evaluation of secondary tiling screens

To validate our initial tiling screen results and evaluate a larger set of CRISPRi systems, we designed a secondary library targeting a subset of 30 essential and 30 non-essential genes from the primary screen (Figure 2a). In addition to dCas9-Kox1 and dCas9-Zim3, we investigated the performance of Kox1 when fused to a methyl-CpG binding protein 2 (Kox1-MeCP2), which has been previously suggested to enhance CRISPRi efficacy (Figure 2b).(21) We also sought to evaluate the impact of KRAB domain placement relative to dCas9, as prior work has found that Zim3 enables stronger repression when fused to the N-terminus rather than the C-terminus.(21) Once again using the Fragmid system to enable direct comparisons of vector architecture, we screened the library with Kox1 and Zim3 on both ends of dCas9.

**Figure 2.**
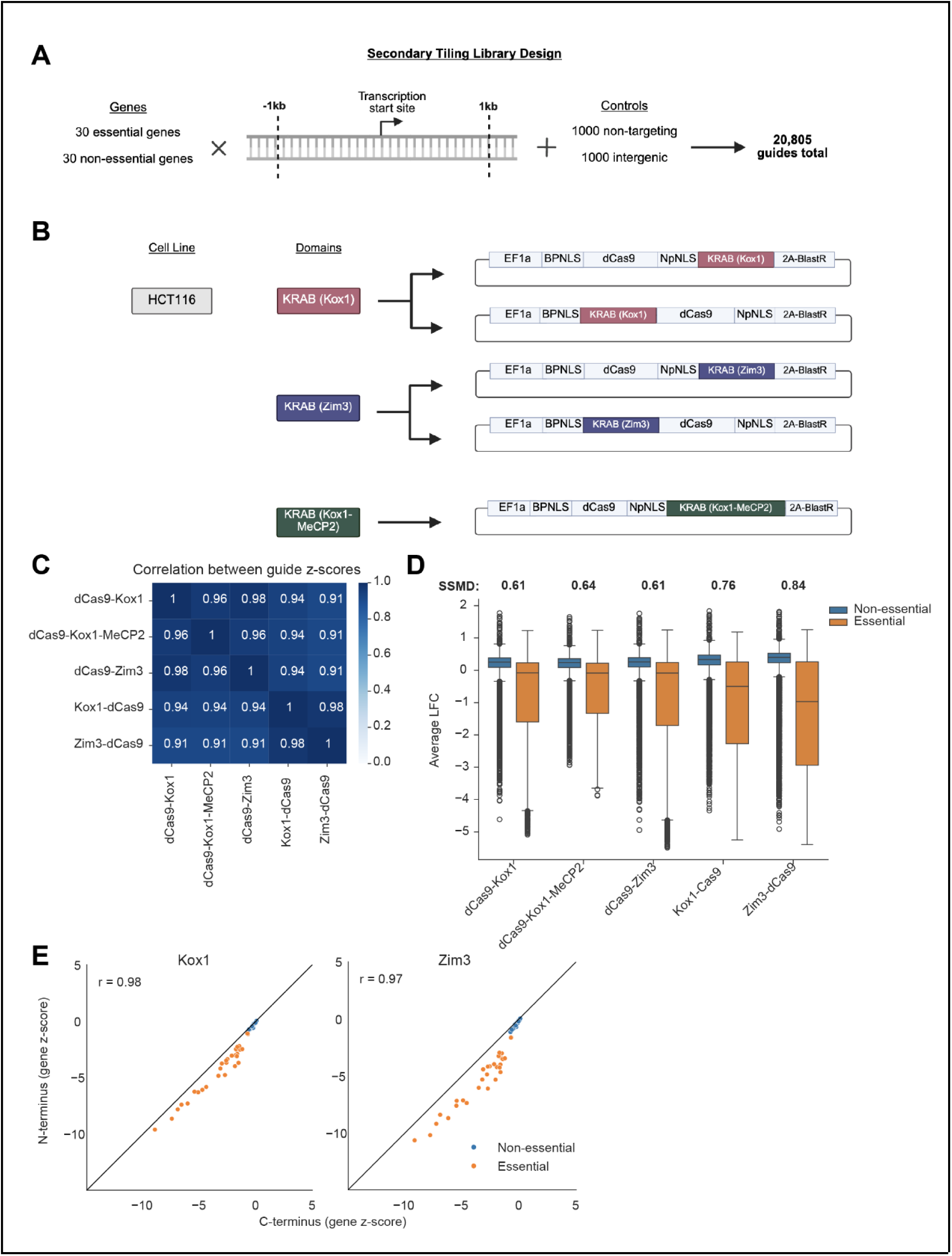
Design and evaluation of secondary tiling screens. (A) Schematic illustrating design and composition of secondary tiling library. (B) Schematic illustrating vector architecture and screening approach of secondary tiling library. (C) Heatmap of Pearson correlation coefficients between guide-level z-scores in each secondary screen performed. Z-scores were computed by averaging log-fold changes across replicates and z-scoring relative to intergenic controls. (D) Boxplot showing average log-fold changes of guides in screens performed with each KRAB domain on each dCas9 terminus. Gene essentiality is indicated by color. Strictly standardized mean difference (SSMD) between non-essential and essential targeting LFCs in each screen is annotated above corresponding boxplots. Boxes show 25th (Q1), 50th (median), and 75th (Q3) percentiles, while whiskers show Q1 - 1.5*IQR and Q3 + 1.5*IQR (where IQR is Q3 - Q1). (E) Scatterplots of gene-level z-scores in screens performed with the Kox1 KRAB domain (left) and the Zim3 KRAB domain (right) tethered to the C-terminus (x-axes) and N-terminus (y-axes) of dCas9. Gene essentiality is indicated by color. Line of equality is shown as a solid black line. Gene-level z-scores were computed by averaging guide-level z-scores of guides targeting the same gene.

The library was screened in replicates in HCT116 cell lines expressing each of the described constructs (Figure 2b). All screens demonstrated strong replicate reproducibility (Pearson R = 0.96-0.97); furthermore, the performance of C-terminal KRAB fusions was highly concordant between the primary and secondary screens at the guide level (Supplementary Figure 2a-b). We observed consistent results across all tested domains and configurations (Figure 2c), as well as similar separation between non-essential and essential targeting sgRNAs among all C-terminal fusions, suggesting that Kox1-MeCP2 does not offer a substantial advantage over Kox1 alone (Figure 2d). N-terminal fusions, however, consistently yielded stronger depletion of essential-targeting guides, with Zim3-dCas9 outperforming Kox1-dCas9 (Figure 2d-e). These results confirm comparable efficacy of multiple repressor domains, demonstrate slightly better performance of N-terminal KRAB fusions relative to C-terminal fusions, and confirm Zim3-dCas9 as an optimal configuration for CRISPRi-mediated repression.

Having established robust CRISPRi efficacy with direct fusion of KRAB domains to dCas9, we next examined if CRISPRi could also be performed using nanobody-mediated recruitment of a repressor domain. We previously used ALFA-nanobody (hereafter abbreviated as ALFA-Nb) recruitment of transactivation domains to successfully activate genes using Cas12a CRISPR activation (CRISPRa) architecture.(29) To test the compatibility of this system with Cas9 CRISPRi, we generated two sets of constructs: guide constructs consisting of a validated CD47 or CD55 targeting guide and a Kox1 or Zim3 KRAB domain fused to either terminus of the ALFA-Nb, and dCas9 constructs consisting of dCas9 linked to 5xALFA tags on either terminus (Supplementary Figure 3a). We tested the resulting 20 construct combinations in HCT116 cells, using guide constructs expressing Zim3-dCas9 as a direct fusion comparison. We found that 5xALFA-dCas9 architecture facilitates greater knockdown than dCas9-5xALFA, consistent with our previous findings with Cas12a (Supplementary Figure 3b-c). In addition, we observed better performance with recruitment of Zim3 than Kox1. The optimal configuration – 5xALFA-dCas9 in combination with Zim3-Nb-ALFA – knocked down CD47 and CD55 expression by 93% and 74% respectively, achieving comparable performance to Zim3-dCas9 (93% and 89% respectively) (Supplementary Figure 3d-e). These results suggest that nanobody recruitment can be used as an alternative to direct KRAB domain fusion to perform Cas9 CRISPRi experiments and potentially facilitate multiplexing.

### Assessing predictive features of CRISPRi on-target activity

We next sought to leverage our primary tiling screen data, in combination with recently updated TSS annotations and chromatin accessibility measurements, to examine predive features of CRISPRi guide efficacy. To increase the statistical power of our analyses, we combined our data with previously published tiling datasets, including another viability screen by Nunez et al. and a ricin susceptibility screen by Gilbert et al. (Figure 3a, Supplementary Data 1).(5, 30) Our analyses were limited to perturbations with expected phenotypes, namely those targeting essential genes(31) in viability screens and genes involved in ricin susceptibility(32) in the ricin screen.

**Figure 3.**
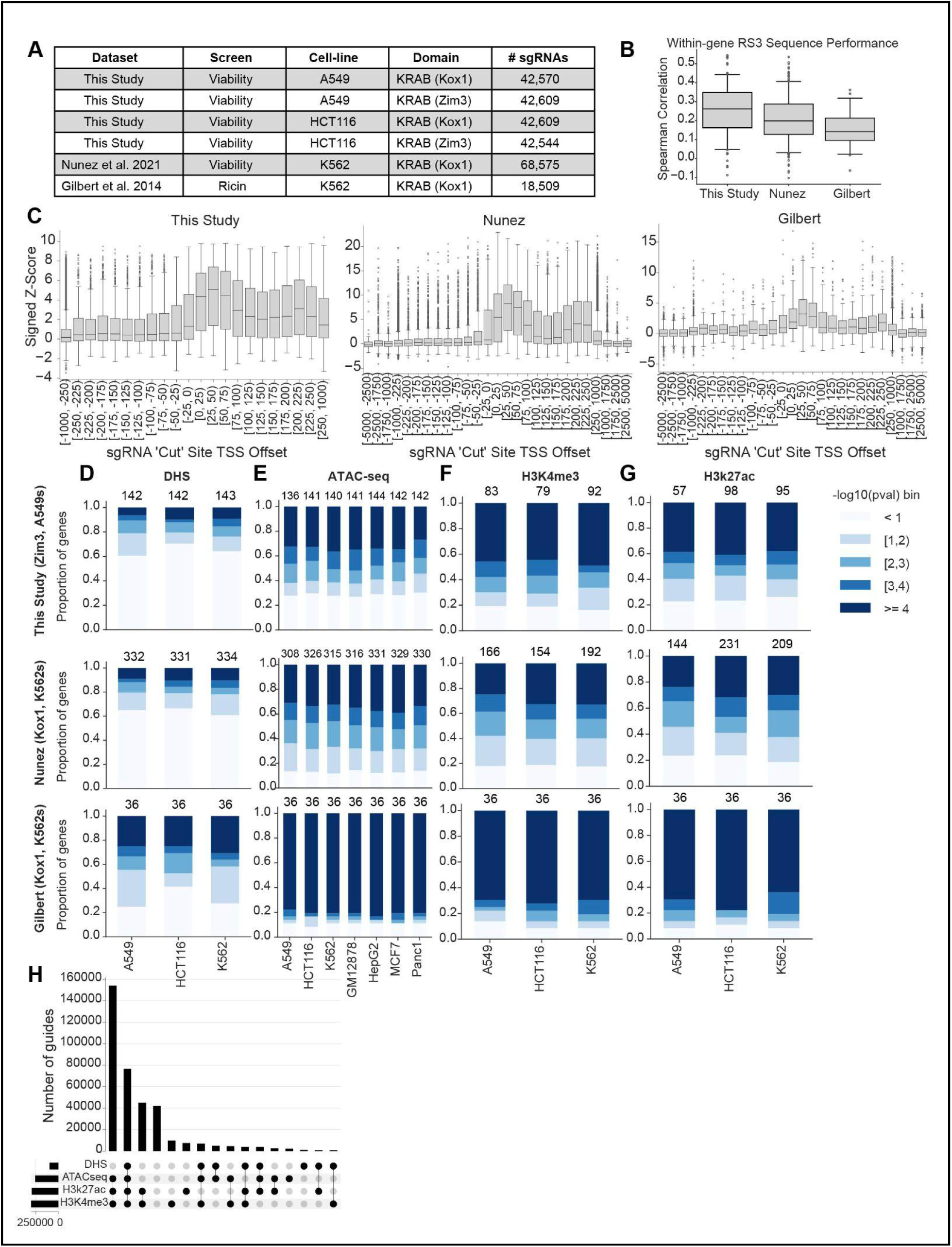
Assessment of predictive features of CRISPRi on-target activity. (A) Composition of curated CRISPRi tiling datasets used for assessment of predictive features after data cleaning (Methods). (B) Spearman correlations between RS3 Sequence score and experimental activity for guides within each dataset. Activity scores were averaged across guides present in all 4 tiling datasets generated in this study. Boxes show 25th (Q1), 50th (median), and 75th (Q3) percentiles, while whiskers show Q1 - 1.5*IQR and Q3 + 1.5*IQR (where IQR is Q3 - Q1). (C) Boxplots of signed z-scores for guides in each dataset (indicated by title), binned by displacement of the sgRNA “cut” site from the MANE Select TSS of its targeted gene. Z-scores were averaged across guides present in all 4 tiling datasets generated in this study, as well as sign-adjusted such that a more positive score corresponds to a more active guide. Boxes show 25th (Q1), 50th (median), and 75th (Q3) percentiles, while whiskers show Q1 - 1.5*IQR and Q3 + 1.5*IQR (where IQR is Q3 - Q1). (D) Proportion of genes targeted in each tiling dataset (indicated on the left) with p-values within the ranges indicated in the color legend. P-values were obtained from one-sided Mann Whitney U tests comparing the z-scores of guides within DHS peaks to those outside of DHS peaks. Bars are stratified by cell line from which DHSs were derived and annotated with the total number of genes for a sufficient number of guides were present in both groups (at least 10 guides within peaks and 10 guides outside of peaks) for a p-value to be calculated. (E) Same as (D) for peaks identified by ATAC-seq. (F) Same as (D) for H3k4me peaks identified by ChIP-seq. (G) Same as (D) for H3k27ac peaks identified by ChIP-seq. (H) Upset plot showing the cardinality of the intersections of guides in curated CRISPRi tiling datasets appearing within peaks identified by each chromatin accessibility dataset type.

We previously developed a sequence-based model for sgRNA on-target efficacy prediction, Rule Set 3 Sequence (RS3 Sequence).(20) While the model was trained on CRISPRko data, we found that it was also predictive of CRISPRi guide performance, suggesting that there are sequence characteristics that similarly influence sgRNA activity regardless of CRISPR modality. Consistent with these findings, we observed a positive correlation between RS3 Sequence score and guide efficacy within our curated tiling datasets (Figure 3b). Likewise, sgRNA position relative to the target TSS is a well-established determinant of CRISPRi efficacy.(6, 7, 15) In all three datasets, guides within the region 0-75bp downstream of the MANE Select TSS showed highest activity, corroborating previous findings (Figure 3c). Conversely, although early studies suggested that CRISPRi sgRNAs targeting the nontemplate strand are more effective than those targeting the template strand(4), we found this difference in activity to be insignificant for the majority of genes (Supplementary Figure 4a).

Horlbeck et al. integrated continuous signals from DNase hypersensitive sites (DHS), MNase-seq, and FAIRE-seq data to predict CRISPRi and CRISPRa guide activity, showing that guides targeting open chromatin regions demonstrate higher efficacy.(6) In our analysis, we chose to evaluate ATAC-seq, as it has now become the method of choice for high-throughput examination of DNA accessibility.(33) Given the known role of nucleosomes in directly obstructing Cas9 access to DNA, we also analyzed histone ChIP-seq datasets.(16) In addition to assessing the utility of these data for predicting CRISPRi sgRNA activity, we sought to determine the extent to which cell line specificity affects their utility.

We first analyzed DHS data, which was used for designing our Dolcetto CRISPRi library. We obtained DHS data collected from A549, HCT116, and K562 cell lines, chosen for their representation in our curated datasets. We then assessed whether sgRNAs within DHS peaks displayed higher activity than those outside of peaks, and found that the difference was not statistically significant for most genes (Figure 3d, Supplementary Figure 4b). We then repeated this assessment with ATAC-seq datasets, including those collected from the three previously analyzed cell lines as well as four other epigenetically well-characterized lines (Panc1, MCF7, GM12878, HepG2). In this case, the majority of genes displayed statistically significant differences in activity between guides targeting within an ATAC peak versus those outside of peaks (Figure 3e, Supplementary Figure 4c). Finally, we repeated this assessment with histone ChIP-seq datasets for two markers for which extensive data are available: H3K4me3 and H3K27ac. sgRNAs within peaks identified by both markers also displayed significantly higher levels of activity than those outside of peaks (Figure 3f-g, Supplementary Figure 4d-e).

We were surprised that these results differed when using DHS versus ATAC-seq datasets; this led us to investigate differences within the underlying data themselves. We observed that approximately half of the guides that appear within ATAC-seq and ChIP-seq peaks are not captured by DHS, suggesting that the latter assay may be less sensitive (Figure 3h). Differences in the peak calling algorithms used by the ENCODE consortium when processing the sequencing results of these assays may also contribute to discrepancies in identified peaks. These results indicate that ATAC-seq and histone ChIP-seq data are likely more valuable than DHS for optimizing CRISPRi sgRNA selection. Notably, with all three types of chromatin accessibility datasets, our observations were consistent across cell lines, suggesting that the insights derived can reasonably be generalized to other human cell lines, although this will vary by gene.

### On-target model development and validation

To integrate these features into an updated on-target model for CRISPRi guide prioritization, we selected 80% of genes in the dataset at random for training, holding out the remaining 20% for testing (Figure 4a). The model was constructed using extreme gradient boosting (XGBoost) to capture the nonlinear relationship between sgRNA offset from the target TSS and activity. Using Shapley Additive Explanation (SHAP) values to evaluate the specific contributions of each feature to the model, we observed that distance from the TSS, RS3 Sequence score, and ATAC peak overlap provided substantially more predictive value than other features (Figure 4b). We thus trained a feature-selected model to predict activity values using only these top three most influential features, which we refer to as Rule Set 3 Interference (RS3i) (Figure 4a). To accommodate the varying numbers of input cell lines when quantifying ATAC peak overlap, we chose to bin values for this feature rather than represent it as a continuous variable.

**Figure 4.**
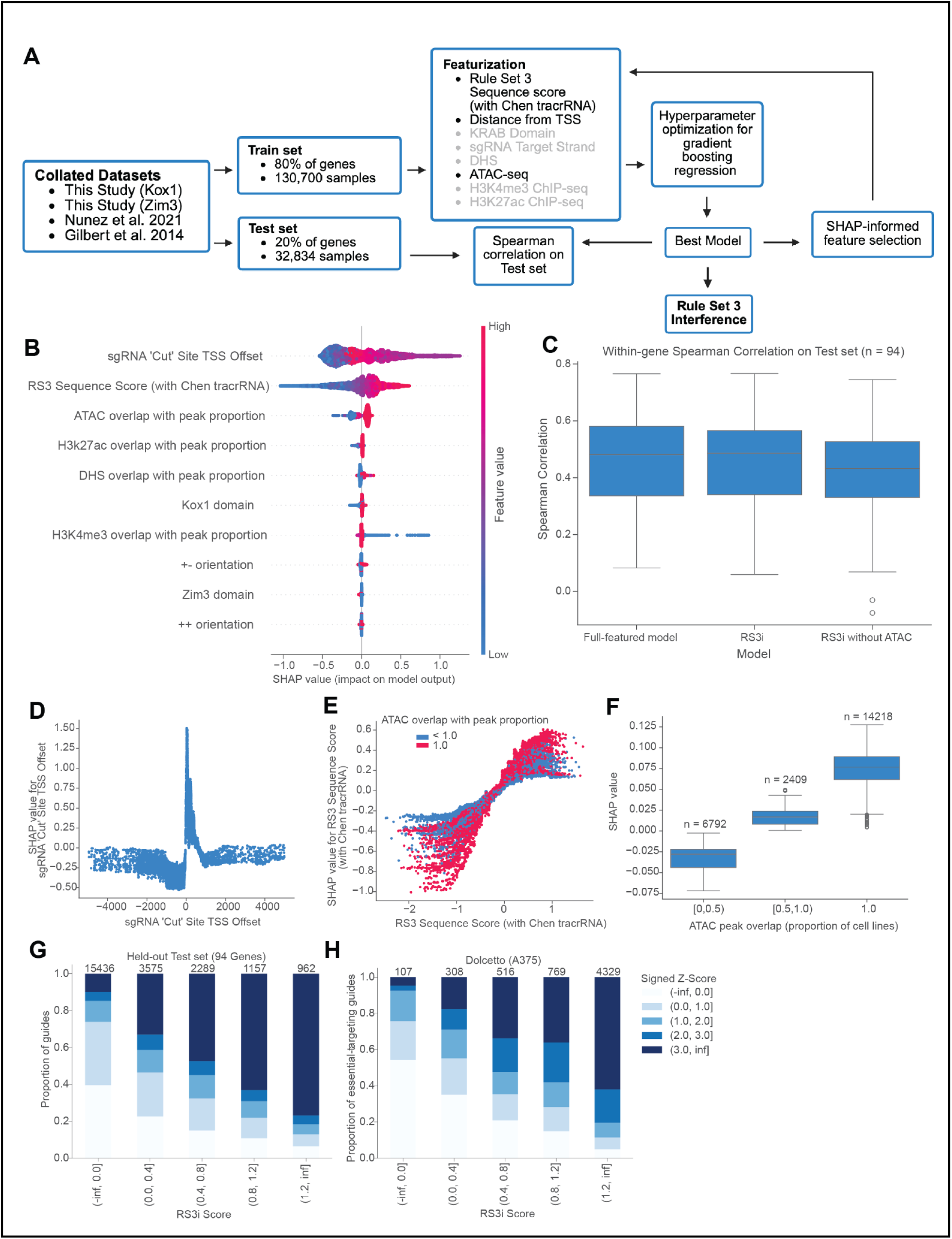
Construction and validation of Rule Set 3 Interference. (A) Schematic illustrating development of Rule Set 3 Interference. Features written in gray were eliminated from the model following analysis of SHAP values. (B) SHAP values for 10 most influential features in the initial, full-featured model. (C) Boxplots showing spearman correlations between activity predicted by each of the models shown on the x axis and experimental activity of all guides targeting each gene in held-out test set. Boxes show 25th (Q1), 50th (median), and 75th (Q3) percentiles, while whiskers show Q1 - 1.5*IQR and Q3 + 1.5*IQR (where IQR is Q3 - Q1). (D) Scatterplot with guides in test set. The x axis shows the target position relative to the MANE Select TSS, and the y axis shows SHAP importance values for the corresponding feature. (E) Same as (D) but for RS3 Sequence score, with points colored by ATAC peak overlap. (F) Boxplot with guides in test set. The x axis shows the (binned) proportion of cell lines in which a guide overlaps with an ATAC peak, as well as the number of guides represented in each boxplot. The y axis shows SHAP importance values for the corresponding feature. Boxes show 25th (Q1), 50th (median), and 75th (Q3) percentiles, while whiskers show Q1 - 1.5*IQR and Q3 + 1.5*IQR (where IQR is Q3 - Q1). (G) Proportion of guides in held-out test set in each range of z-scored log-fold changes (sign-changed such that a more positive z-score indicates a more effective guide), binned by RS3i score. Guide activity was quantified by log-fold changes z-scored relative to intergenic controls. Number of guides within each RS3i score bin is annotated on top of each bar. (H) Same as (G) with essential-targeting guides in Dolcetto screen performed in A375 cells.

To evaluate model performance, we calculated the Spearman correlation between predicted and experimental guide activity for each gene in the held-out test set, a metric that reflects our ultimate goal of selecting the most effective guides per gene. The model demonstrated similar training and testing performance, suggesting that it is not overfit (Supplementary Figure 5a). We also observed comparable performance between the original, full-featured model and Rule Set 3i, justifying the elimination of less important features (Figure 4c, Supplementary Figure 5b). We next examined SHAP partial dependence plots to more closely understand how each selected feature influences model output. SHAP values for TSS offset reflected the same trend observed previously, with values directly upstream of the TSS providing dramatically more impact than others (Figure 4d). Similarly, SHAP values for RS3 Sequence score were positively correlated with the score itself, and guide RNAs overlapping with ATAC-seq peaks found in multiple cell lines tended to have higher SHAP values (Figure 4e-f). We further noticed that RS3 Sequence score is more predictive of activity for guides with high ATAC peak overlap values, suggesting that ATAC peaks shared across many cell lines may contain more highly conserved sequences.

We then asked whether Rule Set 3i could be used when targeting genomes lacking ATAC data. To address these cases, we trained and tested a separate model excluding the ATAC peak overlap feature. Removal of ATAC-seq data reduced model performance, reinforcing the value of the feature (Figure 4c). However, the differences in performance were minor for most genes, suggesting that Rule Set 3i can be valuable for library design in the absence of ATAC-seq data (Supplementary Figure 5b).

To improve our understanding of the model output and its relation to guide efficacy, we investigated the relationship between Rule Set 3i scores and experimental activity. We observed a strong correspondence: of sgRNAs with predicted activity scores above 1.2, 81.6% had signed z-scores > 2, implying that prioritizing guide selection using Rule Set 3i would yield a high percentage of active guides (Figure 4g). Conversely, only 14.7% of those with Rule Set 3i < 0 had signed z-scores > 2. To further validate this observation, we applied the model to a previously published screen using our Dolcetto library.(7) Dividing essential-targeting sgRNAs into the same predicted activity bins, we again observed a positive correlation between Rule Set 3i scores and empirical activity (Figure 4h). 80.4% of guides with predicted activity scores above 1.2 had signed z-scores > 2, while the same was true for only 7.5% of those with Rule Set 3i < 0. These results demonstrate that the model reliably identifies both high-activity and low-activity guides.

### Characterizing seed-driven off-target behavior

The potential for off-target effects in CRISPR screens presents a separate and equally important challenge to address. We recently developed CRISPick Aggregate CFD, a model that leverages our previously established cutting frequency determination (CFD) matrix to estimate cumulative nuclease activity at off-target sites.(34) Mirroring the approach taken to optimize this model, we used depletion of nonessential-targeting and intergenic CRISPRi guides to investigate potential predictors of dCas9 off-target activity (Figure 5a).

**Figure 5.**
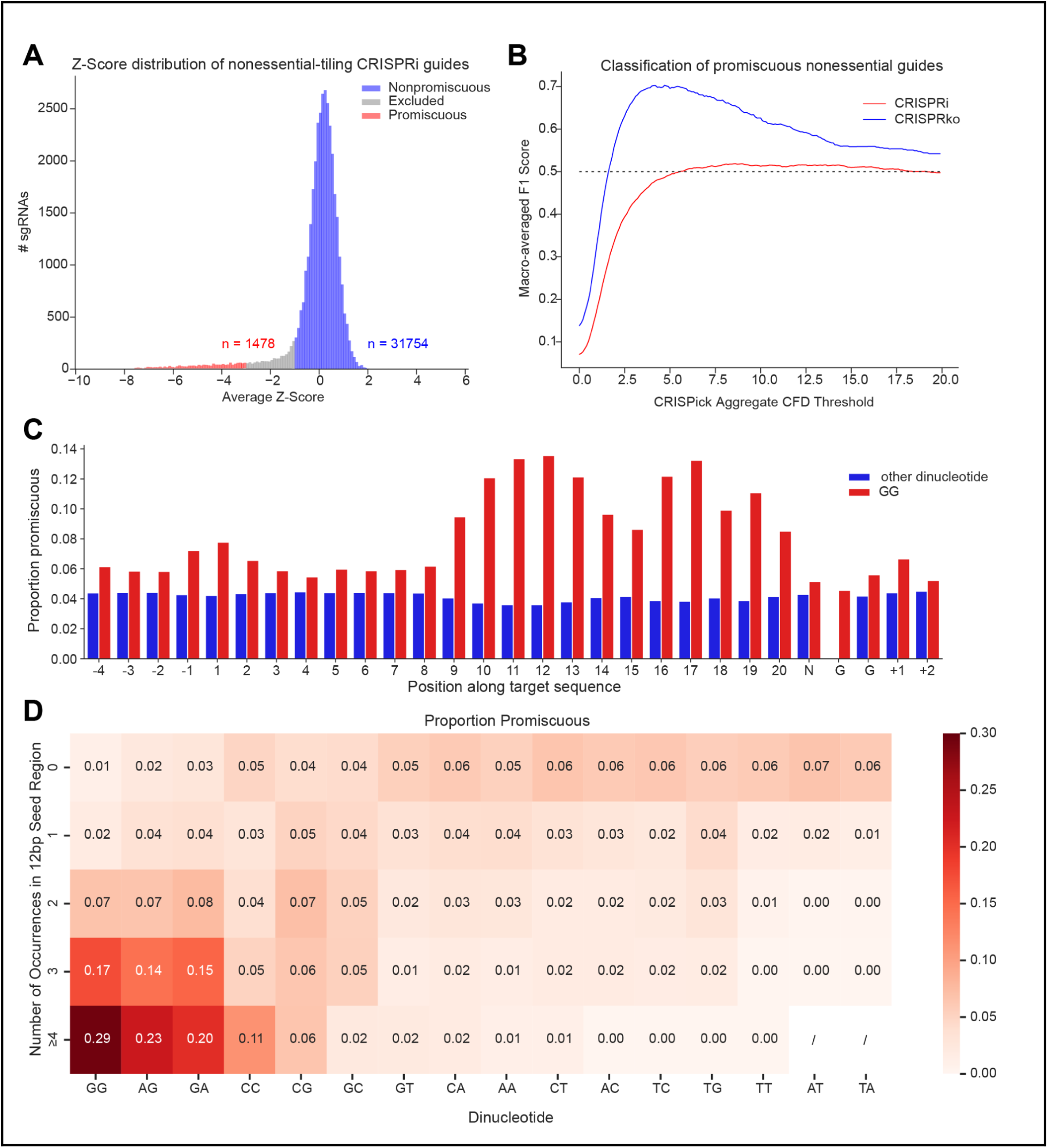
PAM-rich seed sequences promote increased CRISPRi promiscuity. (A) Histogram showing distribution of average z-scored log-fold changes of guides targeting nonessential genes and intergenic controls in primary tiling library, cleaned to exclude those targeting overlapping genes and with low predicted on-target activity (Methods). Z-scores were computed relative to intergenic controls and averaged across the four primary tiling datasets generated in this study. Promiscuous guides (shown in red) were defined as those with a z-score < -3 and nonpromiscuous guides (shown in blue) were defined as those with a z-score > -1. (B) Lineplot showing performance of CRISPick Aggregate CFD as a classifier of promiscuous guides within CRISPRko and CRISPRi nonessential tiling datasets. Promiscuous and nonpromiscuous guides in both datasets were defined as those with z-score < -3 and z-score > -1 respectively. The y axis shows the macro-averaged F1 score of guide classification when guides with CRISPick Aggregate CFD score greater than the threshold shown on the x axis are predicted to be promiscuous. CRISPR mechanism is indicated by color. Horizontal dashed black line indicates performance of a random classifier (0.5). (C) Bar plot showing the proportion of guides that are promiscuous, stratified by presence or lack thereof of a “GG” (indicated by color) at any given position along the target genomic region (indicated on the x axis). (D) Heatmap showing the proportion of guides that are promiscuous as a function of occurrence within the 12bp seed region (indicated on the y axis) of each dinucleotide sequence (indicated on the x axis). Overlapping dinucleotide occurrences are indicated (i.e. GGG contains two GGs). Each colored cell represents a proportion of at least 30 guides. Black slashes indicate dinucleotide-occurrence combinations for which less than 30 guides are present

ChIP-seq experiments performed with dCas9 have demonstrated the importance of the “seed” sequence - broadly defined as the 10-12 base pairs (bp) of the guide RNA most proximal to the protospacer adjacent motif (PAM) - in an intermediate step of Cas9 binding.(24, 25) PAM-proximal mismatches are thus poorly tolerated in both CRISPRko and CRISPRi contexts.(8, 35) Seed-sequence binding is insufficient to trigger nuclease activity, as comprehensive off-target profiling via GUIDE-seq has demonstrated Cas9 is rarely active at off-target sites with multiple mismatches to the guide RNA among the 17 most PAM-proximal nucleotides.(34, 36) In contrast, recent work has demonstrated that mere seed-sequence complementarity can facilitate widespread off-target activity in CRISPRi and CRISPRa contexts, where transient dCas9 binding can impact gene expression.(26) Moreover, given that CRISPRi does not introduce double-stranded breaks, not all off-target effects contribute equally (or at all) to cell viability. We were therefore unsurprised to observe that CRISPick Aggregate CFD, which additively aggregates predicted anti-proliferative effects at a limited set of highly similar off-target sequences, is a poor classifier of promiscuous CRISPRi guides in our dataset (Figure 5b).

Recently, a CRISPRa Perturb-seq screen showed that guide RNAs sharing certain 3-6 bp seed sequences clustered by transcriptomic profile, suggesting the presence of strong seed-driven off-target effects.(27) Many of the identified promiscuity-prone seed sequences were enriched for SpCas9 PAM motifs, and we hypothesized that a similar mechanism could drive dCas9 promiscuity in CRISPRi screens. To investigate this possibility, we first assessed whether the presence of the canonical Cas9 PAM sequence (NGG or simply GG) along the guide RNA sequence was associated with off-target behavior. We found that GG occurrence within the 12 bp seed sequence substantially increased the likelihood that a guide was promiscuous, prompting us to focus subsequent analyses on this region (Figure 5c). We then observed a clear relationship between the count of GGs within the 12bp seed sequence, hereafter referred to as the Seed Score, and probability of off-target activity (Figure 5d). To ascertain the specificity of this trend to PAM sequence occurrence, we assessed the same relationship for every dinucleotide. Notably, we observed the same trend (albeit to a lesser extent) with the dinucleotides AG and GA - noncanonical PAM sequences also known to be recognized by Cas9 - further supporting the hypothesis that PAM-rich seeds promote off-target effects.

To establish a threshold for excluding guides with the potential for off-target activity, we assessed the statistical significance of promiscuous guide overrepresentation among guides with seed sequences enriched for each dinucleotide (Supplementary Figure 6a). We observed the most significant relative abundance of promiscuous guides among those with 3 or more GGs, a trend that generalized to a previous screen conducted with the Dolcetto library (Supplementary Figure 6b). sgRNAs with a Seed Score of 3 or higher capture 52.7% of promiscuous guides within our tiling dataset. Importantly, this group includes only 9% of nonpromiscuous guides, suggesting that imposing this threshold would not unnecessarily eliminate a large fraction of guide options. It is also worth noting that these guides are likely to be suboptimal options given that sgRNAs with Seed Scores of at least 3 display significantly lower on-target activity than others, possibly as a result of increased time spent at off-target loci (Supplementary Figure 6c). These findings suggest that avoiding PAM-rich seed sequences can offer a computationally simple yet reliable strategy for reducing off-target effects in CRISPRi screens.

### Challenges in CRISPRi library design

An additional challenge that complicates the design of compact CRISPRi libraries is the size of the potential target space, specifically because many genes can be expressed from multiple promoters.(37) The MANE Select set includes one transcript per protein-coding locus in the human genome that is most highly supported by the RefSeq and GENCODE catalogs, providing a useful starting point for frequently used TSSs. However, reliance on only these transcripts will fail to capture the use of alternative TSSs in specific cell types or screening contexts. We observed, for example, evidence of ZNF131 expression from an alternative promoter in our tiling dataset: analysis of ZNF131 targeting guides revealed notable depletion near an alternative TSS, rather than directly downstream of the MANE Select TSS (Figure 6a). To quantify the prevalence of alternative TSS usage, we leveraged our curated on-target tiling datasets. Examining the fraction of genes for which we found no evidence of MANE Select transcript downregulation, but instead saw depletion of guides targeting an alternative TSS, we observed false negative rates of 1 - 2.5% across cell lines (Figure 6b). Acknowledging that these data focus on essential genes, which may not be representative of the rest of the genome, this provides assurance that MANE Select annotations can be relied on for the vast majority of targets. Nevertheless, alternative TSS usage presents an important failure mode of CRISPRi.

**Figure 6.**
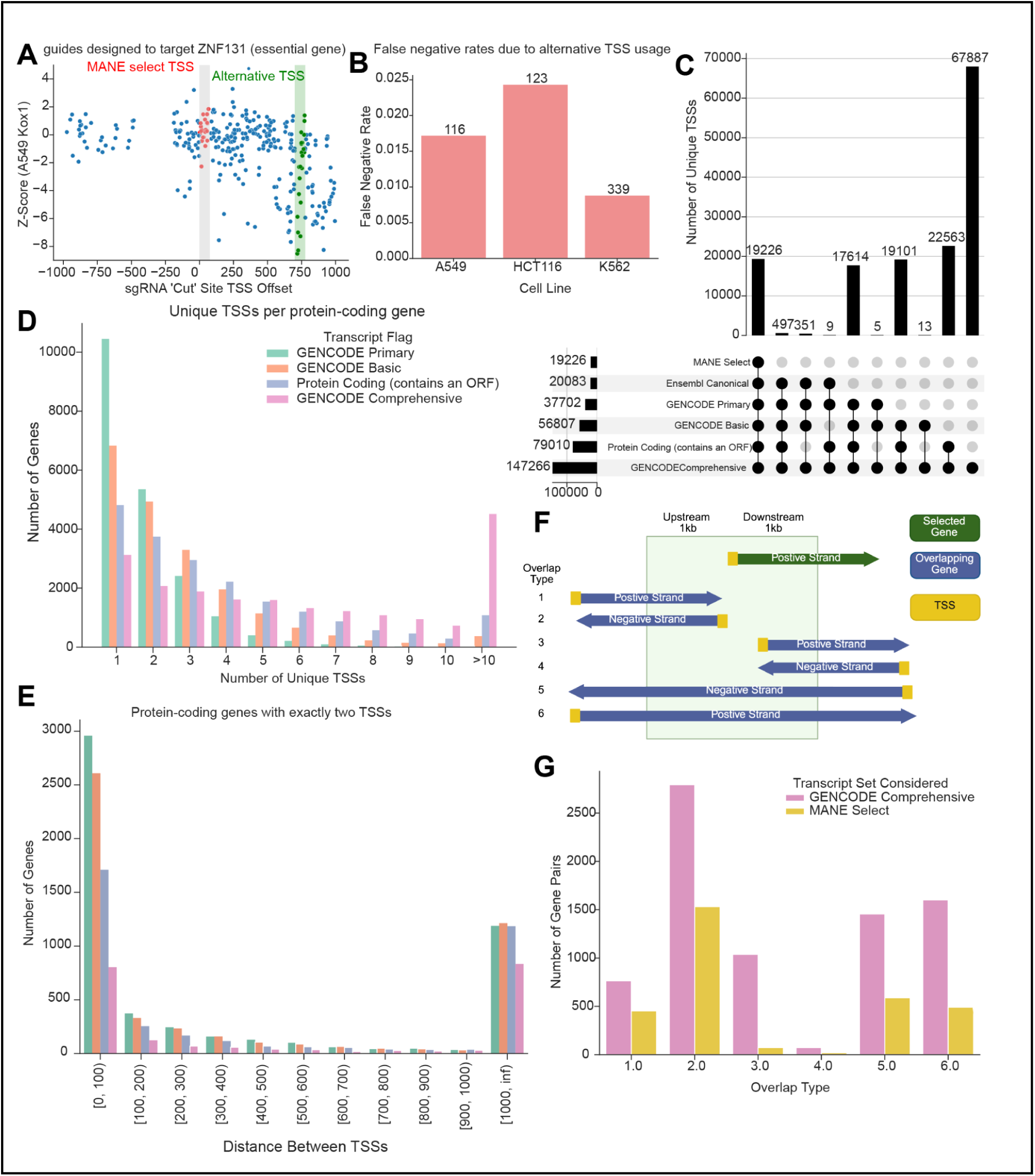
Alternative TSSs and pervasive genomic loci overlap underscore challenges in CRISPRi library design. (A) Scatterplot with guides targeting ZNF131 in the CRISPRi tiling screen performed in A549s with the Kox1 KRAB domain. The x axis shows the target position relative to the MANE Select TSS, and the y axis shows LFC values z-scored relative to intergenic controls. The optimal targeting region (defined as 25-75bp downstream of the TSS) corresponding to the MANE Select TSS is shaded in gray, and guides within this region are labeled in red. The optimal targeting region corresponding to an alternative TSS is shaded in green, and guides within this region are labeled in green. (B) Barplot showing false negative rates due to alternative TSS usage amongst genes in curated on-target datasets. Alternative TSSs were defined as any GENCODE48 TSS not equivalent to the MANE Select TSS. False negatives were determined by selecting the top three guides per TSS according to RS3i score, calculating the average signed z-score per TSS, and defining successful/unsuccessfully downregulated TSSs as those with average signed z-scores above 3 and below 1 respectively (Methods). (C) UpSet plot showing intersection of unique TSSs associated with the MANE Select, Ensembl Canonical, GENCODE Primary, GENCODE Basic, functionally protein-coding, and GENCODE Comprehensive sets transcripts of protein-coding genes. (D) Distribution of the number of unique annotated TSSs present within the GENCODE Primary, GENCODE Basic, functionally protein-coding, and GENCODE Comprehensive transcript sets (indicated by color) for each protein-coding gene. (E) Barplot showing the number of protein-coding genes in the human genome with exactly two unique TSSs in the GENCODE Primary, GENCODE Basic, functionally protein-coding, or GENCODE Comprehensive transcript sets (indicated by same colors used in C), binned by the distance in base pairs between the TSSs. (F) Schematic illustrating potential methods of gene overlap within 1kb. (G) Barplot showing the number of unique protein-coding gene pairs in the human genome that exhibit each type of overlap defined in (E) when all GENCODE transcripts are considered (shown in pink) and when only MANE Select transcripts are considered (shown in yellow).

To characterize the prevalence of alternative TSSs across the human genome, we cataloged all transcripts of protein-coding genes present in GENCODE release 48 (released May 2025), otherwise referred to as the GENCODE Comprehensive set. We first assessed the overlap between unique TSSs present in relevant transcript subsets, including transcripts labeled as functionally protein-coding (containing an open-reading frame), as well as the curated GENCODE Basic, GENCODE Primary, Ensembl Canonical, and MANE Select sets (Figure 6c). We observed that the MANE Select and Ensembl Canonical sets - which by definition include only one transcript per gene - comprise less than a seventh of the entire GENCODE Comprehensive set, with the inclusion of the GENCODE Primary, GENCODE Basic, and protein-coding sets adding approximately 20,000 transcripts each. We then examined the distribution of unique TSS counts per gene. Of the set of 20,120 protein-coding genes, we found that only about half (10,449) possess a single GENCODE Primary TSS. Within the broader GENCODE Basic, protein-coding, and GENCODE Comprehensive transcript sets, this number is reduced to 6,831; 4,812; and 3,125 genes, respectively (Figure 6e). We computed the distance between TSSs for genes with exactly two unique TSSs within each of the aforementioned sets (Figure 6e). This analysis revealed that a substantial number of genes possess TSS pairs that are thousands of base pairs apart, which would require completely separate targeting guides. While the cost of large-scale screens often precludes the possibility of targeting every known TSS, FANTOM CAGE-seq data offer a robust resource for selecting high-confidence secondary TSSs.(17)

Unintentional perturbation of neighboring genes due to overlapping genomic loci presents another critical challenge.(37, 38) Prior work has shown that genes located within 1kb of an essential gene were significantly more likely to arise as a false positive in viability screens.(37) To quantify the scale of this issue across the human genome, we considered six distinct orientations in which genomic overlap may occur (Figure 6f). We focused on the two cases where two separate protein-coding genes are transcribed from a bidirectional promoter, which are most likely to impact CRISPRi sgRNA specificity. When considering the GENCODE Comprehensive set of transcripts, we found that approximately 17% of genes had at least one TSS within 1kb of a TSS belonging to a different gene, similar to previous findings (Figure 6g, Supplementary Data 2).(37) Even when restricting this analysis to only the MANE Select annotations, approximately 8% of genes showed overlap, underscoring the importance of accounting for gene overlap during the design and analysis of CRISPRi screens (Supplementary Data 3). While neighbor-gene targeting is impossible to avoid without sacrificing the on-target activity of selected guides, false positive hit nomination can be mitigated through comprehensive guide annotation and robust downstream analysis methods. These analyses not only highlight challenges specific to CRISPRi library design, but also emphasize the utility of CRISPRko as an orthogonal modality for hit validation.

### Optimized genome-wide Cas9 CRISPRi library design

To design an updated genome-wide Cas9 CRISPRi library, we first collected all Ensembl Canonical transcripts of protein-coding genes listed in GENCODE 48. Seeking to additionally target frequently-used alternative TSSs, we leveraged a recently published analysis that identified a single GENCODE 39 transcript per protein-coding gene supported by the highest average FANTOM5 CAGE-seq read count across all characterized cell types.(39) 85.7% of transcripts identified by this effort - hereafter referred to as the Jaganathan set - are Ensembl Canonical transcripts, lending confidence to the selection strategy. We thus chose to supplement the Ensembl Canonical set with the Jaganathan set.

For each unique TSS in this combined target set, guide RNAs were prioritized in order of RS3i score. To keep the library compact, we selected three guides per target, reasoning that our use of improved guide design principles would enable sensitive detection of hit genes with fewer guides. We first limited guide options to a selective subpool that met strict on-target and off-target criteria in addition to high RS3i scores (Supplementary Data 4). Namely, we prioritized guides with Seed Scores below 3 to minimize the selection of promiscuous guides, as well as preferentially selected those without any off-target sites with a CFD score of 1.0 to avoid target ambiguity. Single nucleotide polymorphisms (SNPs) in certain cell models may also inhibit guide activity; we therefore also preferentially avoided guides located in regions with high sequence variation across the human population as defined by gnomAD variant frequency.(40) We relaxed these criteria one at a time to fulfill a quota of 3 sgRNAs per TSS, allowing for the selection of multi-target guides to ensure that optimal guides were selected for genes with overlapping promoters (Supplementary Data 4).

The resulting library, named Katsano, comprises 62,404 unique targeting sgRNAs corresponding to 20,106 genes. For 1,503 genes, distinct guide sets were chosen to target multiple TSSs (Supplementary Figure 7a). 90.5% of the guides were selected in the first picking round, thereby satisfying our gold standard criteria (Supplementary Data 4). Katsano additionally includes 900 intergenic and 100 non-targeting controls. Intergenic guides were strategically selected to have similar Seed Score profiles to targeting guides, increasing the likelihood that observed differences in phenotype reflect true biological signal rather than off-target effects (Supplementary Figure 7b). As a consequence of updated TSS annotations and refined guide design considerations, a substantial portion of Katsano guides were not included in previous genome-wide Cas9 CRISPRi libraries, despite Katsano’s overall smaller size (Figure 7a). Consistent with our design choices, Katsano guides exhibit higher RS3i scores than Dolcetto and hCRISPRiv2, indicating increased on-target potential (Figure 7b). Similarly, less than 1% of Katsano guides have Seed Scores of 3 or higher, compared to 13.8% and 13.2% of Dolcetto and hCRISPRiv2 guides respectively, indicating reduced off-target potential (Figure 7c).

**Figure 7.**
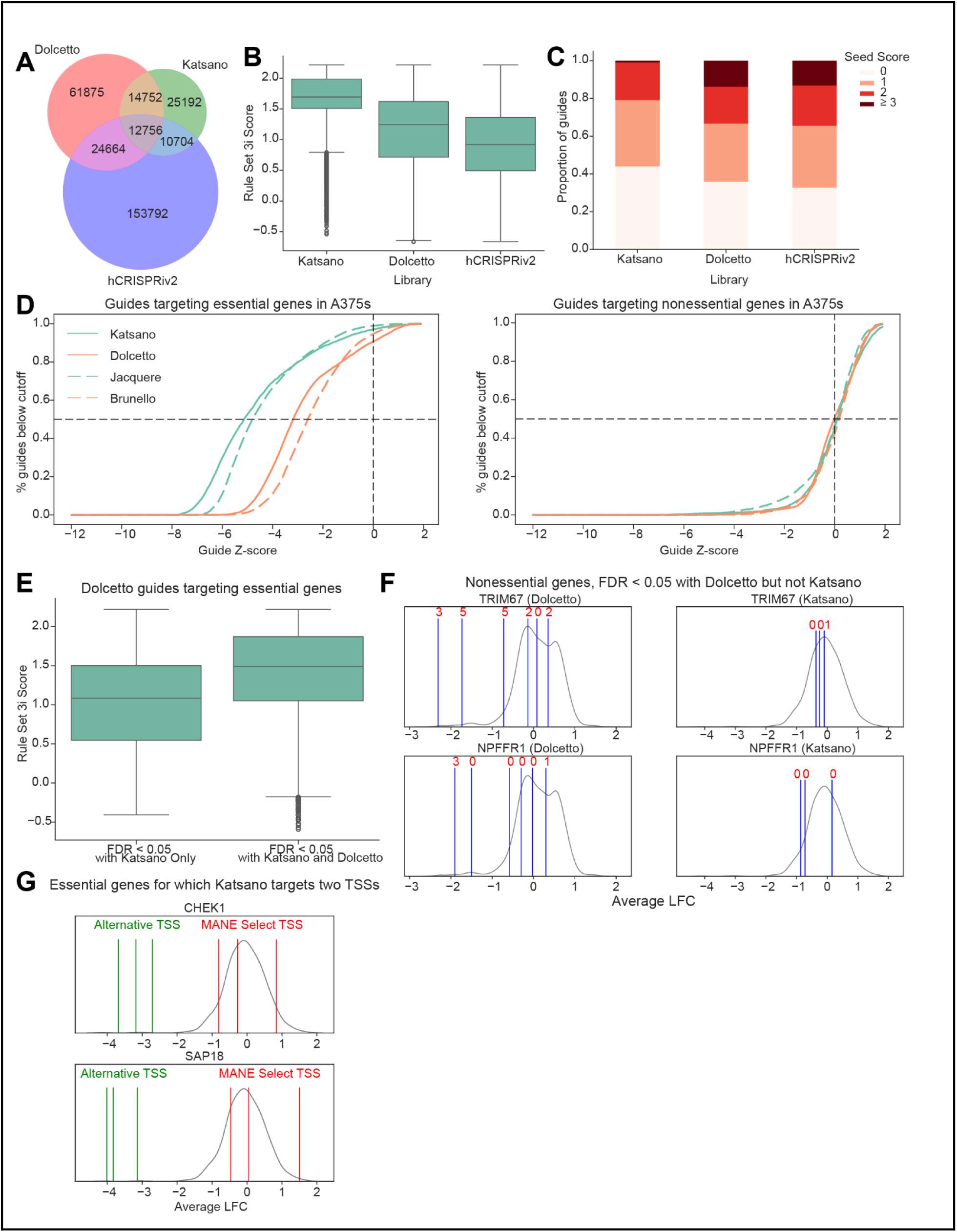
Design and evaluation of an optimized genome-wide Cas9 CRISPRi library. (A) Venn diagram showing overlap between sgRNA sequences in genome-wide Cas9 CRISPRi libraries. Given that hCRISPRiv2 includes guides that are 19 nucleotides long, only the most PAM-proximal 19 bp of the guide sequences are considered in this comparison. (B) Boxplots showing distribution of RS3i scores of targeting guides in genome-wide Cas9 CRISPRi libraries. Boxes show 25th (Q1), 50th (median), and 75th (Q3) percentiles, while whiskers show Q1 - 1.5*IQR and Q3 + 1.5*IQR (where IQR is Q3 - Q1). (C) Proportion of guides in each genome-wide Cas9 CRISPRi library with each Seed Score. (D) Cumulative z-score distribution of guides targeting essential (left) and nonessential (right) genes in genome-wide CRISPR screens in A375 cells. Library with which the screen was performed are indicated by color and line style. Z-scores are calculated relative to intergenic controls. Horizontal and vertical dashed black lines indicate a cumulative proportion of 0.5 and a z-score of 0 respectively. (E) Boxplots showing distribution of RS3i scores of Dolcetto guides targeting essential genes identified as hits in A375s with Katsano and/or Dolcetto, as indicated on the x axis. Boxes show 25th (Q1), 50th (median), and 75th (Q3) percentiles, while whiskers show Q1 - 1.5*IQR and Q3 + 1.5*IQR (where IQR is Q3 - Q1). (F) Plots showing guide-level log-fold changes of nonessential genes identified as hits in A375s with Dolcetto but not Katsano. Distributions of intergenic control log-fold changes are shown in gray, and blue lines indicate log-fold changes of targeting guides. Gene and library are indicated in subplot titles. (G) Plots showing guide-level log-fold changes of essential genes identified as hits (FDR < 0.05) in A375s through targeting of an alternative TSS rather than the MANE Select TSS. Distributions of intergenic control log-fold changes are shown in gray, red lines indicate log-fold changes of guides targeting the MANE Select TSS, and green lines indicate log-fold changes of guides targeting the alternative TSS.

### Screening performance of Katsano

To evaluate Katsano, we conducted viability screens in A375 and K562 cell lines expressing Zim3-dCas9. Our choice of cell lines enables direct comparisons to Dolcetto, our recently developed Cas9 CRISPRko library Jacquere, and a multitude of published CRISPRi datasets.(7, 34, 38) We observed good reproducibility between replicates in both cell lines (Pearson R of 0.66 and 0.89 in A375s and K562s respectively), as well as strong separation between guides targeting nonessential and essential genes (Supplementary Figure 7c). As expected, MAPK signaling pathway genes BRAF and MAP2K1 showed selective vulnerability in BRAF-mutant A375 melanoma cells, while downregulation of the BCR-ABL oncogene caused depletion exclusively in K562 leukemia cells (Supplementary Figure 7d). Lineage-defining transcription factors SOX10 and GATA1 likewise depleted selectively in A375s and K562s respectively. The Katsano library shows remarkably greater on-target activity than Dolcetto, as evidenced by depletion of essential genes, without any meaningful change in depletion of non-essential genes, indicating low off-target activity (Figure 7d). Additionally, Katsano performs similarly to our recent Cas9 knockout library, Jacquere, highlighting the value of these two libraries as orthogonal and equally effective screening resources (Figure 7d).(34)

At a false discovery rate (FDR) of 5%, Katsano recovered all but 63 essential genes as significant hits in A375 cells, whereas Dolcetto failed to recover 95, indicating that Katsano achieves a lower false negative rate at the same level of statistical confidence with half the number of guides per target. These essential genes missed by Dolcetto were targeted by guides with lower predicted on-target activity than those recovered as hits by both libraries, demonstrating the value of prioritizing guides by RS3i score (Figure 7e). Similarly, the value of avoiding guides with high Seed Scores is illustrated by nonessential genes falsely identified as hits by Dolcetto but not Katsano (Figure 7f). Finally, Katsano also offers the potential to uncover true positives through downregulation of alternative TSSs, as exemplified by the recovery of essential genes CHEK1 and SAP18 (Figure 7g). Together, these results demonstrate strong screening performance of Katsano and reinforce the value of the guide selection strategies employed for its design.

## DISCUSSION

Here we performed large-scale CRISPRi tiling screens with which we compared performance of several previously characterized CRISPRi systems. We showed that KRAB domains from both Kox1 and Zim3 enable stronger repression when tethered to the N-terminus of dCas9 rather than the C-terminus, with the Zim3 N-terminal fusion demonstrating highest efficacy. We combined these data with other CRISPRi tiling data to investigate predictive features of CRISPRi on-target activity and ultimately developed a new on-target model, which integrates sgRNA sequence features, distance from transcription start site, and presence within open chromatin regions defined by ATAC-seq peaks. We verified and further refined previous observations that guide RNAs with PAM-rich seed sequences are more likely than others to exhibit off-target behavior, suggesting a simple approach for avoiding promiscuous guides. We leveraged these newly developed design rules to construct a genome-wide Cas9 CRISPRi library, named Katsano, which targets a comprehensive set of frequently expressed transcripts and demonstrates superior screening performance compared to existing libraries.

Effective CRISPRi sgRNAs are constrained to a narrow window proximal to the TSS, substantially limiting active guide options per target relative to CRISPRko. Nevertheless, we demonstrate here that with data-driven selection of sgRNAs with high on-target potential, CRISPRi achieves comparable efficacy to CRISPRko, offering a useful alternative for examining copy number amplified loci, contexts in which double-stranded breaks are poorly tolerated, or studies for which incomplete suppression is desired.(7, 37) However, targeting bidirectional promoters with CRISPRi has the potential to yield false positives when a dependency overlaps with a non-hit gene, presenting an important limitation. Here, CRISPRko may provide a useful orthogonal modality by which to validate observed phenotypes.

Another important limitation of CRISPRi arises from alternative TSS usage. Katsano is designed to be cell-line agnostic and thus aims to target TSSs that are experimentally supported across many cell types. However, alternative TSSs in specific screening models and contexts is inevitable. For custom screens performed in specific biological models of interest, it may be worth characterizing the model’s transcriptome beforehand to ensure comprehensive coverage of expressed isoforms. This is especially prudent given the negligible one-time cost of characterizing a valuable model with bulk RNA-sequencing relative to that of numerous large-scale genetic screens in that same model.

We anticipate continual improvement in CRISPRi on-target activity predictions as transcript annotations are further refined. In particular, the advent of deep learning models for predicting transcription initiation enables our guide selection strategies to be extended to organisms with less well-characterized genomes.(41) In the future, given that CRISPR activation (CRISPRa) is mechanistically similar to and shares guide efficacy determinants with CRISPRi, we plan to employ similar approaches to investigate and re-inform our Cas9 CRISPRa library design methods.(6, 7) Together, the optimizations presented here will enable CRISPRi to be more effectively leveraged for functional interrogation across a multitude of screening contexts.

## MATERIALS AND METHODS

Key reagents and resources are reported in Supplementary Data 5.

### Vectors

#### Modular vector design

All pF plasmids were made by gene synthesis into the EcoRV site of pUC57-Kan (Genscript) as part of the Fragmid vector design kit (Addgene 1000000237).

#### Modular vector destination vector pre-digest

Destination vectors were pre-digested using BbsI and NEBuffer 2.0 (New England Biolabs) at 37°C for 2 h. Linearized destination vectors were gel purified using 0.7% agarose gels and extracted with the Monarch DNA Gel Extraction Kit (New England Biolabs), before further purification by isopropanol precipitation.

#### Modular vector Golden Gate assembly

The pF vectors were diluted to 10 nM in sterile water and cloned into a pre-digested destination vector via Golden Gate cloning. Each reaction contained 3 μL BbsI (New England Biolabs), 1.25 μL T4 ligase (New England Biolabs), 3 μL of 10x T4 ligase buffer (New England Biolabs), 75 ng destination vector, and a 1:1 molar ratio of fragments:destination vector. Reactions were carried out under the following thermocycler conditions: (1) 37°C for 5 min; (2) 16°C for 5 min; (3) go to (1), x100; (4) 37°C for 30 min; (5) 65°C for 20 min. The Golden Gate product was treated with Exonuclease V (New England Biolabs) at 37°C for 30 min before enzyme inactivation with the addition of EDTA to 11 mM. Per reaction, 10 μL of product was transformed into Stbl3 chemically competent E. coli (Invitrogen) via heat shock, and grown at 37°C for 16 h on agar with 100 μg/mL carbenicillin. Colonies were picked and grown at 37°C for 16 h in 5 mL Luria-Bertani (LB) broth with 100 μg/mL carbenicillin. Plasmid DNA (pDNA) was prepared (QIAprep Spin Miniprep Kit, Qiagen). Purified plasmids were verified by restriction enzyme digest and whole plasmid sequencing through Plasmidsaurus.

### Cell lines and culture

All cells regularly tested negative for mycoplasma contamination and were maintained in the absence of antibiotics except during screens, flow cytometry-based experiments, and lentivirus production, during which media was supplemented with 1% penicillin-streptomycin. Cells were passaged every 2-3 days to maintain exponential growth; for adherent cells, this meant seeding at ∼20% confluence at passaging when they achieved ∼90% confluence; for K562 cells, a suspension cell line, cells were seeded at 3 x 10^4 - 1.2 x 10^5 cells per mL and passaged at ∼5.3 x 10^5 6.5 x 10^5 cells/mL. Cells were kept in a humidity-controlled 37°C incubator with 5.0% CO2. Media conditions and doses of polybrene, puromycin, and blasticidin were as follows, unless otherwise noted:

A549: DMEM + 10% fetal bovine serum (FBS); 1ug/mL; 1.5ug/mL; 5ug/mL.

HCT116: McCoy’s 5A + 10% FBS; 4ug/mL; 3ug/mL; 16ug/mL.

A375: RPMI 1640 + 10% FBS; 1ug/mL; 1ug/mL; 5ug/mL

K562: RPMI 1640 + 10% FBS; 1ug/mL; 1ug/mL; 8ug/mL

HEK293T: DMEM + heat-inactivated FBS; N/A; N/A; N/A.

### Library production

Oligonucleotide pools for CP1948, CP2084, and CP2240 were synthesized by TWIST. BsmBI recognition sites were appended to each sgRNA sequence along with the appropriate forward and reverse overhang sequences for cloning into the sgRNA expression plasmids, as well as primer sites to allow differential amplification of subsets from the same synthesis pool.

Primers were used to amplify individual subpools using 25 μL 2x NEBnext PCR master mix (New England Biolabs), 2 μL of oligonucleotide pool (∼30-300 ng), 5 μL of primer mix at a final concentration of 0.5 μM, and water to a final volume of 50 µL. PCR cycling conditions: (1) 98°C for 1 min; (2) 98°C for 30 sec; (3) 53°C for 30 s; (3) 72°C for 30 s; (4) go to step 2, x6-14 depending on library size; (5) 72°C for 5 min. The resulting amplicons were PCR-purified (Qiagen) and cloned into their respective library vector via Golden Gate cloning with Esp3I (Fisher Scientific) and T7 ligase (Epizyme) under the following thermocycler conditions: (1) 37°C for 5 min; (2) 20°C for 5 min; (3) go to step 1, x100; (4) 37°C for 30 min; (5) 65°C for 10 min. The ligated product was isopropanol precipitated and electroporated into Stbl4 electrocompetent cells (Invitrogen) and grown at 37°C for 16 h on agar with 100 μg/mL carbenicillin. Colonies were scraped and plasmid DNA (pDNA) was prepared (HiSpeed Plasmid Maxi, Qiagen). To confirm library representation and distribution, the pDNA was sequenced by Illumina MiSeq.

### Lentivirus production

For small-scale Cas virus production for tiling screens, the following procedure was used: 24 h before transfection, HEK293T cells were seeded in 6-well dishes at a density of 1.5 x 10^6 cells per well in 2 mL of DMEM +10% heat-inactivated FBS. Transfection was performed using TransIT-LT1 (Mirus) transfection reagent according to the manufacturer’s protocol. Briefly, one solution of Opti-MEM (Corning, 66.75 uL) and LT1 (8.25 uL) was combined with a DNA mixture of the packaging plasmid pCMV_VSVG (Addgene 8454, 125 ng), psPAX2 (Addgene 12260, 1250 ng), and the transfer vector (e.g., the library pool, 1250 ng). The solutions were incubated at room temperature for 20–30 min, during which time media was changed on the HEK293T cells. After this incubation, the transfection mixture was added dropwise to the surface of the HEK293T cells, and the plates were centrifuged at 1000 g for 30 min at room temperature. Following centrifugation, plates were transferred to a 37 C incubator for 6–8 h, after which the media was removed and replaced with DMEM +10% FBS media supplemented with 1% BSA. Virus was harvested 36 h after this media change.

For small-scale Cas virus production for Katsano screens, the following procedure was used: 24 h before transfection, HEK293T cells were seeded in 6-well dishes at a density of 1.0 x 10^6 cells per well in 1mL DMEM +10% heat-inactivated FBS. Transfection was performed using TransIT-LT1 (Mirus) transfection reagent according to the manufacturer’s protocol. Briefly, one solution of Opti-MEM (Gibco, 324 uL) and LT1 (17 uL) was combined with a DNA mixture of the packaging plasmid pCMV_VSVG (280 ng), psPAX2 (2800 ng), and the transfer vector (312.5 ng). The solutions were incubated at room temperature for 20–30 min. After this incubation, the transfection mixture was added dropwise to the surface of the HEK293T cells and the plates were transferred to a 37 C incubator for 6–8 h. Following incubation, the media was removed and replaced with 2.5mL DMEM +10% FBS media supplemented with 1% BSA. Virus was harvested 36 h after this media change.

A larger-scale procedure was used for pooled library virus production. 24 h before transfection, 18×10^6 HEK293T cells were seeded in a 175 cm2 tissue culture flask and the transfection was performed the same as for small-scale production using 6 mL of Opti-MEM, 305uL of LT1, and a DNA mixture of pCMV_VSVG (5 ug), psPAX2 (50 ug), and 40 ug of the transfer vector. Flasks were transferred to a 37°C incubator for 6–8 h; after this, the media was aspirated and replaced with BSA-supplemented media. Virus was harvested 36 h after this media change.

### Derivation of stable cell lines

In order to establish the dCas9 expressing cell lines for tiling screens, A549 and HCT116 cells were transduced with pRDB_181 (dCas9-KRAB(Kox1)); pRDB_182 (dCas9-KRAB(Zim3)); pRDB_331 (KRAB(Kox1)-dCas9); pRDB_332 (KRAB(Zim3)-dCas9; and pRDB_333 (dCas9-KRAB-MeCP2). For Katsano screens, A375 and K562 cells were transduced with pRDB_332. For transduction, cells were seeded in a 12 well plate at 1.5e6-3e6 cells per well in 2mL of medium with polybrene (medium and polybrene concentration listed above for each cell type) along with the appropriate amount of virus. Cells were centrifuged at 2000rpm for 2 hours at 30C. Post centrifugation, 2mL of medium without polybrene was added to each well. The cells were transferred to a 37C incubator for 6-8 hours. After incubation, cells were collected, pooling all replicate wells, and replated. Successfully transduced cells were selected with blasticidin for a minimum of 2 weeks. Cells were taken off blasticidin at least one passage before transduction with the libraries.

### Pooled screens

For all pooled screens, cells were transduced in 2 biological replicates with a lentiviral library. Transductions were performed at a low multiplicity of infection (MOI ∼0.35), using enough cells to achieve a representation of at least 500 transduced cells per sgRNA assuming a 20% - 40% transduction efficiency. The transduction protocol was the same as listed above. Puromycin was added 2 days post-transduction and cells were passaged on puromycin 2-3 times to ensure complete removal of non-transduced cells. When selection was complete, cells were passaged every 2-3 days for an additional 2 weeks to allow sgRNAs to enrich or deplete; cell counts were taken at each passage to monitor growth. At the conclusion of each screen, cells were pelleted by centrifugation, resuspended in PBS, and frozen promptly for genomic DNA isolation.

### Genomic DNA isolation, PCR, and sequencing

Genomic DNA (gDNA) was isolated using the KingFisher Flex Purification System with the Mag-Bind Blood & Tissue DNA HDQ Kit (Omega Bio-Tek), per the manufacturer’s instructions. The gDNA concentrations were measured by Qubit. For samples where genomic DNA was limited, gDNA was purified prior to PCR using the Zymo OneStep PCR Inhibitor Removal Kit (Zymo), per the manufacturer’s instructions. For PCR amplification, gDNA was divided into 100 µL reactions such that each well had at most 10 µg of gDNA. Plasmid DNA (pDNA) was also included at a maximum of 100 pg per well. Each well of a 96-well PCR plate contained 1.5 µL of Titanium Taq (Takara), 10 µL of Titanium Taq buffer, 8 µL of dNTPs, 5 µL of DMSO, 0.5 µL of P5 primer at 100 µM stock, 10 µL of P7 primer at 5µM stock, and sterile water added to 100 µL. PCR cycling conditions were as follows: (1) 95°C for 1 min; (2) 94°C for 30 s, (3) 52°C for 30 s, (4) 72°C for 30 s, (5) go to step 1, x28; (6) 72°C for 10 min. PCR products were purified with Agencourt AMPure XP SPRI beads according to manufacturer’s instructions (Beckman Coulter, A63880). Samples were sequenced using Illumina technology (either MiSeq, HiSeq, or NovaSeq) with a 5% spike-in of PhiX.

Guide sequences were extracted from sequencing reads by running the PoolQ tool (https://portals.broadinstitute.org/gpp/public/software/poolq) with the search prefix “CACCG”. Reads were counted by alignment to a reference file of all possible guide RNAs present in the library. Reads were then assigned to a condition (e.g., a well on the PCR plate) on the basis of the 8 nt index included in the P7 primer.

### Flow cytometry assays with nanobody vectors

HCT116 cells were transduced with virus for each of the dCas9-containing vectors separately; 2 days after transduction, cells were selected with blasticidin for 14 days. Blasticidin was removed for one passage and cells were subsequently transduced with virus for the guide-containing vectors. 3 days after transduction, cells were selected with puromycin for 5 days. Following selection (7 days post-transduction), cells were visualized by flow cytometry on a CytoFLEX S Sampler. To prepare samples for visualization, cells were stained with a fluorophore-conjugated antibody targeting the respective cell surface marker gene, diluted 1:100 for 20-30 minutes on ice.

CD47: APC anti-human CD47 antibody (Biolegend, 323124)

CD55: FITC anti-human CD55 antibody (Biolegend, 311305)

Cells were washed with PBS two times to remove residual antibody and were resuspended in flow buffer (PBS, 2% FBS, 5 μM EDTA). Fluorophore signal was measured in the respective channel (APC-A or FITC-A). Flow cytometry data were analyzed using FlowJo (v.10). Gates were set such that ∼1% of cells score as APC-positive or FITC-positive in the control condition.

### Screen analysis

Following deconvolution, the resulting matrix of read counts was first normalized to reads per million within each condition by the following formula: reads per guide RNA / total reads per condition x 10^6^. Reads per million were then log2-transformed by first adding one to all values, which is necessary in order to take the log for sgRNAs with zero reads. Prior to further analysis of each screen, we filtered out guides for which the log-normalized reads per million within the pDNA were > 3 standard deviations below the pDNA mean. We then calculated log2-fold changes (LFCs) by subtracting log-normalized reads for each condition from normalized pDNA reads (for viability screens). We assessed the Pearson correlation between LFCs of biological replicates, confirming reproducibility before averaging values across replicates. Guide-level z-scores were then obtained by z-scoring LFCs relative to the mean and standard deviation of negative control LFCs. For in-house screens, only guides targeting a singular intergenic site were used as negative controls. For external datasets, any guide labeled as a negative control in the original dataset was used as a negative control.

Guides were mapped to all Ensembl genes that they target in accordance with GENCODE48 annotations, rather than only their intended targets. For CRISPRi, this was defined as location within [-50,300] base pairs of the MANE Select TSS. Z-scores of all guides targeting the same gene were then aggregated using Stouffer’s method. To determine gene-level statistical significance, we generated an empirical null distribution by constructing 1,000 “pseudogenes” by randomly sampling negative controls in the library (with replacement). The number of guides sampled per pseudogene was set to the mode of the number of guides per gene in the library. Z-scores of guides associated with the same pseudogene were similarly aggregated using Stouffer’s method and the resulting distribution of pseudogene z-scores was smoothed using a kernel density estimation. One-sided gene-level p-values were calculated as the proportion of this distribution that fell below each gene z-score, approximating the probability of achieving a higher level of depletion by random chance. False discovery rates were then estimated from p-values using the Benjamini-Hochberg procedure for multiple testing. Previously established common essential and nonessential genes were used as positive and negative controls respectively for evaluating library performance.(31, 42)

### Design of tiling libraries

The primary tiling library targeted the same set of 201 essential genes and 198 nonessential genes targeted in our previously published Rule Set 3 Validation Cas9 CRISPRko tiling library.(20) All potential sgRNAs tiling within 1,000 base pairs of the MANE Select TSS of these genes were identified using CRISPick. The library was then filtered to exclude guides with polyT sequences, BsmBI sites, or greater than 10,000 off-target sites; this removed 19,790 sequences. 1,000 controls targeting intergenic regions and 1,000 non-targeting controls were additionally included, resulting in a total of 108,574 sgRNAs.

The secondary tiling library targeted a subset of 30 essential genes and 30 nonessential genes targeted by the primary tiling library. Selected essential genes were required to have a gene effect score less than or equal to -1 in A549 and HCT116 cell lines according to DepMap, as well as absence of any neighboring TSSs within 1,000 base pairs of their own TSS. We then ranked the essential genes that met these criteria by gene-level z-score (calculated using Stouffer’s method), averaged ranks across the four primary screens conducted, and selected 10 genes from the top, middle, and bottom of the list respectively. The resulting set of 30 genes were targeted by an average of 314 guides per gene. 30 nonessential genes with no neighboring TSSs within 1,000 base pairs were selected with the goal of maintaining comparable representation of essential and nonessential targeting guides within the library. The secondary library included the same set of sgRNAs tiling each gene as within the primary library, as well as the same sets of intergenic and non-targeting controls, comprising a total of 20,805 sgRNAs.

### SSMD Calculation

The strictly standardized mean difference (SSMD) between the log-fold changes of non-essential and essential targeting sgRNAs was calculated with the following formula: 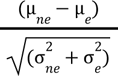, in which “ne” stands for nonessential and “e” stands for essential.

### External data cleaning

Raw read counts for the CRISPRi tiling viability screen published by Nunez et al. were downloaded from Supplementary Table 4 of the the source publication.(30) We ran CRISPick with the genes targeted in this screen as input and default settings for Cas9 CRISPRko to determine the chromosome each gene lies on. Three genes (EFTUD1, QARS, SARS) were unable to be resolved by CRISPick and were therefore excluded from further analyses. Using the retrieved chromosome numbers and the DNA strand and coordinate indicated in the “id” column of the read count table, we used LiftOver to map guide coordinates from the hg19 genome build to the hg38 genome build. There were 3863 coordinates which are no longer present in hg38; those guides were therefore discarded. We then used pysam (version 0.23.3) to retrieve sgRNA context sequences corresponding to each guide. We removed any sequences which did not match the guide sequence indicated in the supplementary data (507 guides) or did not include a PAM at the expected location within the context sequence (4 guides).

Raw read counts from the CRISPRi tiling ricin screen published by Gilbert et al. were obtained via personal communication. These data included guide identifiers that indicated the target gene, sgRNA sequence length, and genomic coordinate of the PAM, but lacked the sgRNA sequences themselves. However, in a later study, the same library was used to perform a ricin screen with nuclease-active Cas9, and these data did include sgRNA sequences, along with the same guide identifiers.(16) We downloaded these CRISPRko screening data from the source publication to retrieve the guide sequences. sgRNA context sequences (defined as 30 nucleotide sequences that include the PAM + 3 nucleotides following the spacer sequence) were obtained by using biopython to query the genome using the supplied PAM coordinates. Since sgRNAs of 25 base pairs or longer are consequently not fully captured by a 30 nucleotide context sequence, they were excluded from further analyses.

Dolcetto, Jacquere, and hCRISPRiv2 screening data were downloaded from their source publications.(6, 7, 34) Guide-gene mappings were obtained for Dolcetto and Jacquere using an internal mapping tool. Since hCRISPRiv2 includes guide sequences that are 19 bp long, we used CRISPick to obtain 20bp guide sequences for this library. Context sequences, distance from the MANE Select TSS of the target gene, and ATAC-seq peak overlap for each guide RNA were obtained from CRISPick. RS3 Sequence scores for all context sequences were calculated using rs3 (version 0.0.15), assuming use of the Chen tracrRNA.

### Cleaning and featurization of on-target datasets

Any gene with at least one TSS located within 1kb of at least one other TSS belonging to another gene was excluded from the combined dataset used for feature selection and model training, with the intent of preventing unintentional perturbation of neighboring gene expression from confounding observed phenotypes.

MANE Select TSS coordinates of all genes were obtained by running CRISPick. The coordinate used to define the target location of each sgRNA was the “cut” position of the sgRNA as defined by CRISPick, which is the position of the sgRNA sequence 3 base pairs from the “N” of the PAM. For datasets generated in this study, the sgRNA target strand was determined from CRISPick; for external datasets, the sgRNA target strand was obtained from supplementary data from the source publications. RS3 Sequence scores for all context sequences were calculated using rs3 (version 0.0.15), assuming use of the Chen tracrRNA.

All analyzed chromatin accessibility datasets were downloaded from ENCODE. We used bed narrowPeak files with peaks called at an FDR cutoff of 5% using DHS data, and bigBed files containing pseudo replicated peaks derived from ATAC-seq and histone ChIP-seq data, all aligned to the GRCh38 reference genome. 2 DHS replicates for each cell line were manually combined and merged using bedtools. When multiple ATAC-seq or ChIP-seq files were available, the file derived from the largest number of isogenic replicates was selected. A guide RNA was defined as within a peak if its “cut” site fell within any peak (inclusive of the start and end positions of the peak) within a particular dataset. One-sided Mann-Whitney U tests were conducted to test the alternative hypothesis that guides within peaks display higher activity than guides outside of peaks.

### On-target modeling

The activity of each guide within the curated on-target datasets was quantified by z-scored log-fold change relative to negative controls, and sign adjusted such that a more positive z-score reflected a more active guide. In order to normalize for the essentiality of each gene, all z-scores were transformed with a second z-score transformation relative to guides targeting the same gene in the same dataset. Following that, double z-scores were averaged if they corresponded to the same context sequence and repressor domain present in multiple datasets. For each sgRNA, we included as features the Rule Set 3 Sequence score (assuming use of the Chen tracrRNA), the displacement of the sgRNA “cut” site from the MANE Select TSS of its target gene, the strands of the target gene and sgRNA, and the repressor domain the sgRNA was screened with. In addition to including sgRNA “cut” site TSS offset as a continuous feature, we categorized TSS offset values into 25bp bins to observe whether decreasing the resolution of this feature would prevent overfitting. We quantified chromatin accessibility of the sgRNA target site by the proportion of cell lines in which it falls within a peak, and included 4 features describing the overlap of each guide with peaks identified by ATAC-seq, DNase-seq, H3K4me3 ChIP-seq, and H3k27ac ChIP-seq respectively.

We selected 80% of the data for training (375 genes, 98,011 unique context sequences, 130,700 samples) and held out the remaining 20% for testing (94 genes, 23,419 unique context sequences, 32,834 samples), ensuring that all sgRNAs targeting a gene were either in the train set or the test set. We also confirmed that each of the original dataset sources (this study, Nunez et al., and Gilbert et al.) were approximately equally represented in the train and test sets. We then used XGBoost (version 2.1.1) to optimize a gradient boosting regression model on the selected training set. We tuned the L1 and L2 regularization terms (between 10^-3^ and 10 on a log_10_ scale), subsampling fraction (between 0.4 and 1), feature subsampling fraction per tree (between 0.4 and 1), maximum tree depth (between 2 and 10), learning rate (between 0.001 and 0.3), and number of estimators (between 500 and 5000) using a tree-structured Parzen estimator from Optuna (version 3.6.1) to optimize the hyperparameter search. Each hyperparameter set was evaluated by performing 4-fold cross validation stratified by gene (using GroupKFold from scikit-learn version 1.3.0) and calculating the mean RMSE across test sets in each fold, and the mean RMSE was minimized over 100 trials. Feature importance was examined using the shap package (version 0.46.0).

Following feature selection, double z-scores corresponding to the same context sequence screened with separate repressor domains were averaged, resulting in smaller training and testing sets including 98,011 and 23,419 samples respectively. Rule Set 3i and Rule Set 3i (Without ATAC-seq) were then developed using the same hyperparameter tuning pipeline outlined above, with the exception that the number of estimators was restricted between 3 and 100. Rule Set 3i has been incorporated into CRISPick (https://portals.broadinstitute.org/gppx/crispick/public).

### Rule Set 3i validation on Dolcetto

Dolcetto guides were subsetted to those targeting previously defined essential genes.(31) Guides targeting overlapping genes were excluded, as were any context sequences present within the Rule Set 3i training set.

### Nonessential tiling data cleaning

Any gene with at least one TSS located within 1kb of at least one other TSS belonging to another gene was excluded from the set of nonessential-targeting guides used to analyze CRISPRi off-target activity, again with the intent of removing neighbor gene knockdown as a potential confounding variable. We also removed guides with a RS3 Sequence score below -0.5, recognizing that sgRNAs with low propensity for on-target activity are also unlikely to display evidence of off-target activity. Reasoning that guides intended to target a singular intergenic site should function similarly to those targeting nonessential genes, we included intergenic controls in the dataset as well. Guide activity was defined as the average z-score of the guide across all four tiling datasets generated in this study. We defined promiscuous guides as those with a z-score below -3, and nonpromiscuous guides as those with a z-score above -1, excluding those with intermediate z-scores to reduce ambiguity.

### Comparing CRISPick Aggregate CFD performance for CRISPRko and CRISPRi

CRISPick Aggregate CFD training data and corresponding context sequences were downloaded from the source publication.(34) To maintain consistency with Aggregate CFD training, we removed sgRNAs with RS3 Sequence scores below 0.2 from both the CRISPRi and CRISPRko datasets. We also excluded genes with an average guide-level z-score below -2, interpreting this as evidence that these genes do not behave as nonessential in the particular cell line they were screened in. Promiscuous and nonpromiscuous guides were equivalently defined in both datasets as those with z-scores below -3 and above -1 respectively. Aggregate CFD scores were obtained by providing CRISPick with raw sequence inputs (consisting of the context sequences of interest concatenated together), selecting CRISPRko as the CRISPR mechanism, and SpyoCas9 as the enzyme.

### Seed Score validation on Dolcetto

Dolcetto guides were subsetted to those targeting previously defined nonessential genes.(42) To remain consistent with the criteria used to clean the nonessential tiling dataset used to establish the Seed Score, we excluded guides targeting overlapping genes, those with RS3 Sequence scores below -0.5, and those targeting genes with an average z-score below -2.

### Calculating false negative rates due to alternative TSS usage

We extracted the set transcripts of protein-coding genes from the GENCODE48 GFF3 file including all transcript annotations on the primary assembly (gencode.v48.primary_assembly.annotation.gff3), found on the GENCODE Release 48 downloads page (https://www.gencodegenes.org/human/release_48.html). After subsetting to transcripts with a “gene_type” attribute of “protein_coding”, presence within the MANE Select, GENCODE Primary, and GENCODE Basic sets of transcripts was determined from the “tag” attribute, and only transcripts with a “transcript_type” attribute of “protein_coding” were considered to contain an open reading frame (ORF). Every annotated TSS for each gene within the curated on-target datasets (matched based on gene symbol) was considered as a potential alternative TSS for that particular gene (if not the MANE Select transcript). We calculated Rule Set 3i scores for each sgRNA with respect to every MANE Select and alternative TSS, and then selected the top three guides per TSS (after excluding those with a Seed Score of 3 or higher on the basis of potential for off-target activity). Signed z-scores for each unique guide were averaged across screens performed in the same cell line, after which they were averaged across guides targeting the same TSS to define TSS-level z-scores. We defined successfully downregulated TSSs as those with an average z-score above 3, and unsuccessfully downregulated TSSs as those with an average z-score below 1, excluding those with intermediate z-scores to reduce ambiguity. A “false negative due to alternative TSS usage” was then defined as a gene for which targeting the MANE Select TSS was unsuccessful but targeting at least one alternative TSS was successful.

### Characterizing scale of gene overlap genome-wide

Transcript annotations for the entire human genome were downloaded from Ensembl Biomart on January 3rd, 2024 (representative of GENCODE45 annotations). The exported dataset was then subsetted to transcripts of genes with a “Gene type” of “protein_coding” before determining pairs of overlapping genes. When considering the GENCODE Comprehensive set of transcripts, two genes were considered to overlap if any pairs of corresponding transcripts overlapped in one of the ways illustrated in Figure 6f. MANE Select transcript IDs were obtained from GENCODE48 annotations.

### Katsano Design

We extracted the set of transcripts of protein-coding genes from the GENCODE48 primary assembly GFF3 file, and identified Ensembl Canonical transcripts as those having an “Ensembl_canonical” tag. The Jaganathan set of GENCODE39 transcripts was downloaded from the source publication (supplementary table 3), and transcript IDs present in GENCODE48 were additionally included in the Katsano target set.(39) For Jaganathan set transcripts no longer present in GENCODE48, we included the closest existing transcript of the same gene symbol or Ensembl gene ID, reasoning that the CAGE-seq evidence with which the transcript was selected indicated the existence of a relevant TSS nearby. Finally, we remove transcripts lacking a corresponding protein ID (39 transcripts), considering them irrelevant for our purpose of functional investigation. These transcripts have a “transcript_type” of “retained_intron” or “protein_coding_CDS_not_defined”. The final Katsano target set included 20,120 Ensembl Canonical transcripts, supplemented by 1927 additional transcripts from the Jaganathan set. As a result, Katsano targets two TSSs with distinct guide sets for several genes (Supplementary Figure 7a). As a consequence of gene nomenclature updates, there are also two genes for which we included three target transcripts: MKKS and AGAP6. The Illumina set included three MKKS transcripts, each previously listed under a different gene ID but unified under the same gene ID in GENCODE48. For AGAP6, we included the Ensembl Canonical and Illumina-selected transcripts. We additionally included ENST00000311652, which was included in the Illumina-selected set as a fusion transcript corresponding to TIMM23B-AGAP6 but is now associated with AGAP6 only.

We obtained the TSS coordinates (including the chromosome, strand, and 1-based genomic coordinate of the TSS) of each target transcript and supplied them as inputs to CRISPick with the following parameters:

- Reference genome: Ensembl v.114
- Mechanism: CRISPRi
- Enzyme: SpyoCas9
- On-Target Scorer: RS3i-Chen2013+ATAC
- Off-Target Scorer: Seed Score v1
- Library Mode (100 non-targeting, 900 intergenic, 0 positive controls/preselects)
- Quota: 4
- Target Local Mode

Four coordinates, all on unlocalized contigs, were not recognized by CRISPick and therefore not targeted.

Under default settings, CRISPick identifies all potential guides within [-50,300] bp of each provided TSS coordinate, excluding those with BsmBI recognition sequences (to enable the use of BsmBI for Golden Gate vector assembly) or containing a run of four or more thymines (to avoid premature termination of Pol III transcription). Potential guide options were then ranked according to the criteria indicated in Supplementary Data 4, with gnomAD variable sites defined as those with a variant frequency greater than 5% across the total human population or greater than 12.5% within the African American population subgroup. Even after relaxing most of these criteria completely, we fully excluded any guides with greater than 10 off-target sites with a CFD score of 1.0, reasoning that they provide minimal insight into the function of any individual gene. Although CRISPick Aggregate CFD was not informative for predicting CRISPRi guide promiscuity, we observed a significant association between the presence of more than 10 CFD=1.0 off-target sites and elevated guide promiscuity (one-sided Fisher Exact test p-value = 0.098), justifying exclusion of these guides. Consequently, 11 additional genes (the majority of which are from the FAM90A and USP17L1 families), are not targeted by Katsano.

Intergenic controls were sampled from a list of 10,000 options with a minimum RS3 Sequence score of 1.0 to ensure comparable on-target activity to targeting guides, and selected to have similar Seed Score distributions to targeting guides, ensuring comparable levels of off-target activity. Although we chose to subset the official Katsano library to the top three guides per target, we provide designs including four guides per target (Supplementary Data 6).

### Data visualization

Figures were created with Python3, FlowJo (v.10), and GraphPad Prism (v.10). Schematics were created with BioRender.com.

## Data availability

Data presented in this manuscript are available on GitHub along with all code used for analyses and figures presented: https://github.com/ssrikant2/Katsano-Manuscript.

The GitHub has also been archived on Zenodo (https://doi.org/10.5281/zenodo.18773405).

## ACKNOWLEDGEMENTS

We thank the Genetic Perturbation Platform, especially Thomas Green and Mark Tomko for developing the infrastructure to enable the analyses described in this manuscript and integrating the novel library design scheme into our web tool, CRISPick. We additionally thank Emanuel Gonçalves for a close reading of the manuscript, Dany Gould for data collation, and Antonino Napoleone and Jessie Li for experimental assistance.

## AUTHOR CONTRIBUTIONS

Conceptualization: SS, FZ, JGD; Formal Analysis: SS, FZ; Investigation: SS, FZ, LMD, STS, EGK, DG, GOL, GR, SM; Visualization: SS, FZ, ATU; Writing - Original Draft: SS, FZ, STS; Writing - Review and Editing: LMD, JGD

## SUPPLEMENTARY FIGURES

**Supplementary Figure 1.**
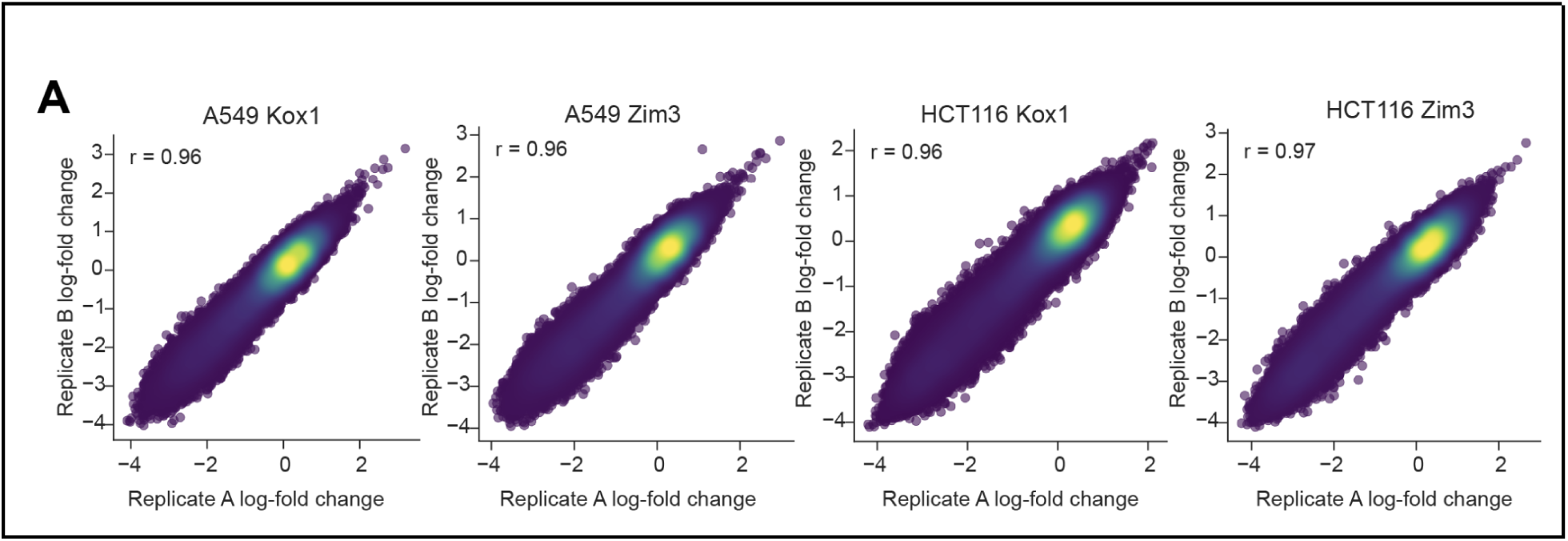
Evaluation of large-scale CRISPRi tiling screens. (A) Scatterplots comparing log-fold changes of biological replicates for each screen. Pearson correlation coefficient displayed.

**Supplementary Figure 2.**
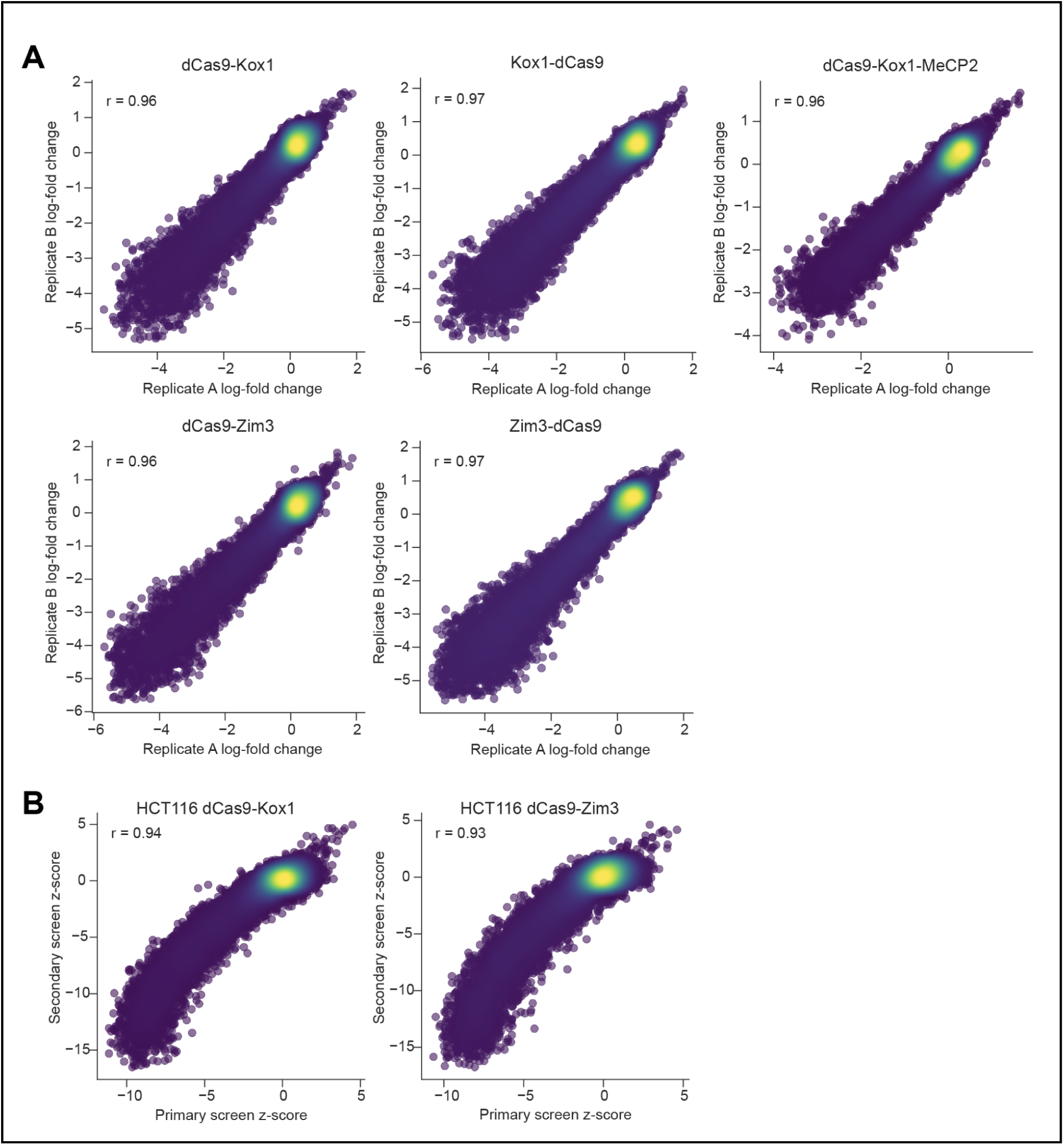
Evaluation of secondary tiling screens. (A) Scatterplots comparing log-fold changes of biological replicates for each screen. Pearson correlation coefficient displayed. (B) Scatterplots comparing guide level z-scores in the primary (x-axes) and secondary (y-axes) screens performed with the Kox1 KRAB domain (left) and the Zim3 KRAB domain (right). Replicated screens were performed with dCas9-KRAB constructs in HCT116 cells.

**Supplementary Figure 3.**
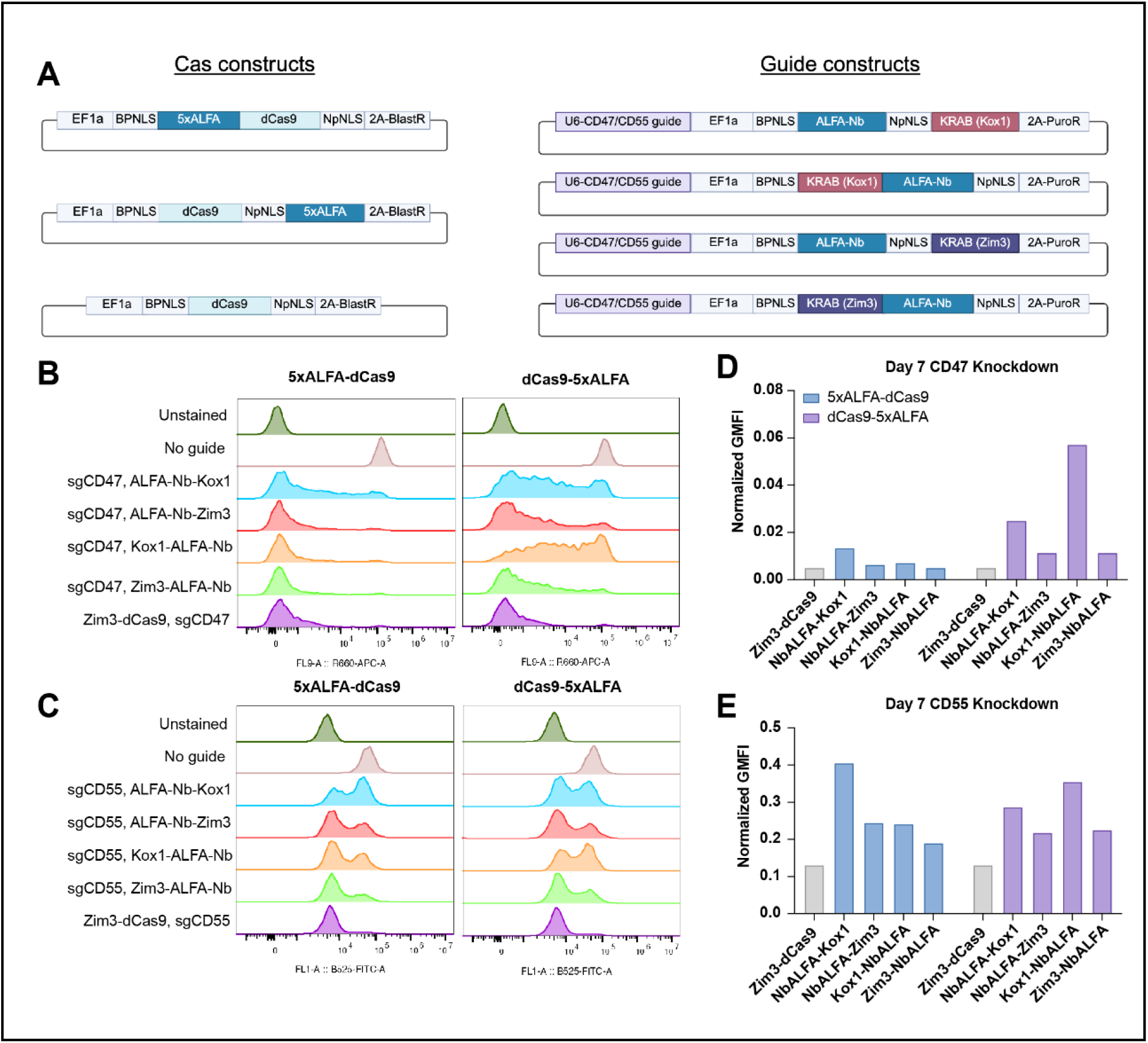
Nanobody-based recruitment of KRAB domains. (A) Schematic illustrating vectors used to test ALFA-nanobody recruitment of KRAB domains. (B) Histograms show expression levels of CD47 in HCT116 cells expressing the constructs denoted on the y axis and titles of the plots, 7 days post transduction. (C) Same as (B) for CD55 expression levels. (D) Barplot showing normalized geometric mean fluorescence intensity (GMFI) of CD47 expression in HCT116 cells expressing the guide constructs denoted on the x axis, 7 days post transduction. The Cas construct used is denoted by color. GMFI values were normalized to cells expressing dCas9 along with a non-targeting guide. (E) Same as (D) for CD55 expression.

**Supplementary Figure 4.**
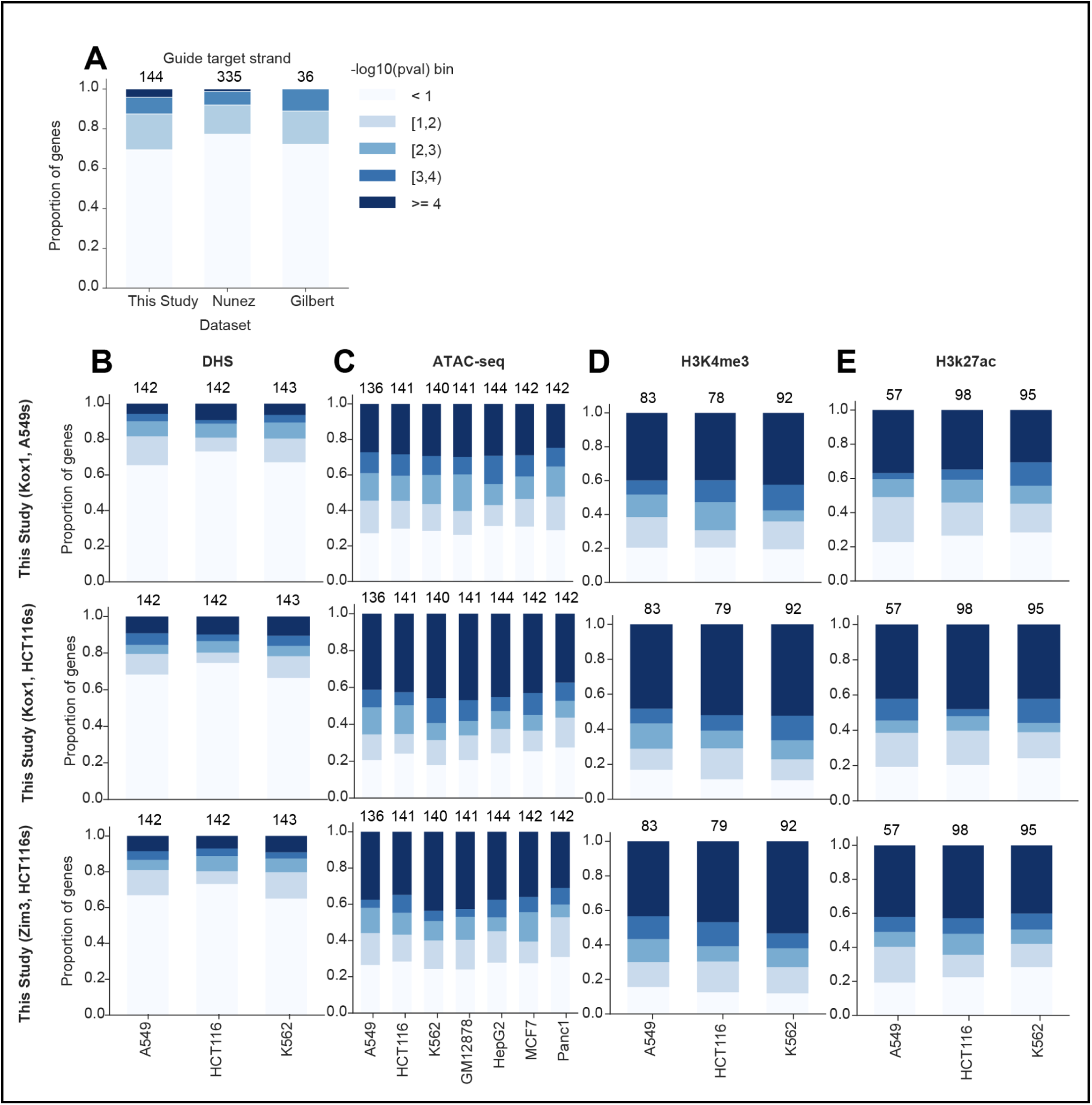
Assessment of predictive features of CRISPRi on-target activity. (A) Proportion of genes targeted in each tiling dataset (indicated on x axis) with p-values within the ranges indicated in the color legend. P-values were obtained from two-sided Mann Whitney U tests comparing the z-scores of guides targeting the template strand vs the nontemplate strand for each gene. Z-scores were averaged across guides present in all 4 tiling datasets generated in this study. Bars are annotated with the total number of genes for which a sufficient number of guides were present in both groups (at least 10 targeting the template strand and at least 10 targeting the nontemplate strand) for a p-value to be calculated. (B) Proportion of genes targeted in each tiling dataset (indicated on the left) with p-values within the ranges indicated in the color legend. P-values were obtained from one-sided Mann Whitney U tests comparing the experimental activity scores of guides within DHS peaks to those outside of DHS peaks.

**Supplementary Figure 5.**
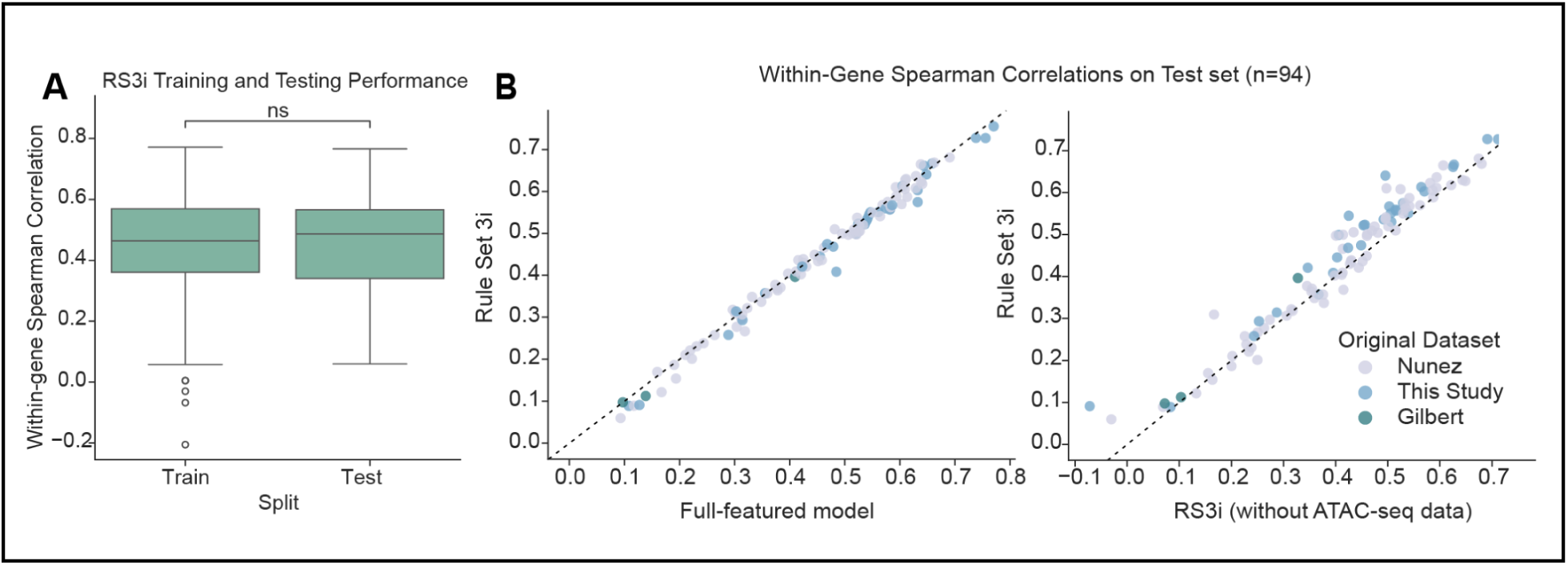
**Validation of Rule Set 3 Interference** (A) Boxplots showing spearman correlations between Rule Set 3i score and experimental activity of all guides targeting each gene in train and test sets. Significance bar indicates the result of a two-sided Mann Whitney U test. Boxes show 25th (Q1), 50th (median), and 75th (Q3) percentiles, while whiskers show Q1 - 1.5*IQR and Q3 + 1.5*IQR (where IQR is Q3 - Q1). (B) Scatterplots of spearman correlations between predicted and experimental activity for all guides targeting each gene in held-out test set. Performance Rule Set 3i is shown on the y axis, while performance of the full-featured model (left) or Rule Set 3i without ATAC-seq (right) is shown on the x axis. The original dataset from which data was taken is indicated by color. Line of equality is shown as a dashed black line.

**Supplementary Figure 6.**
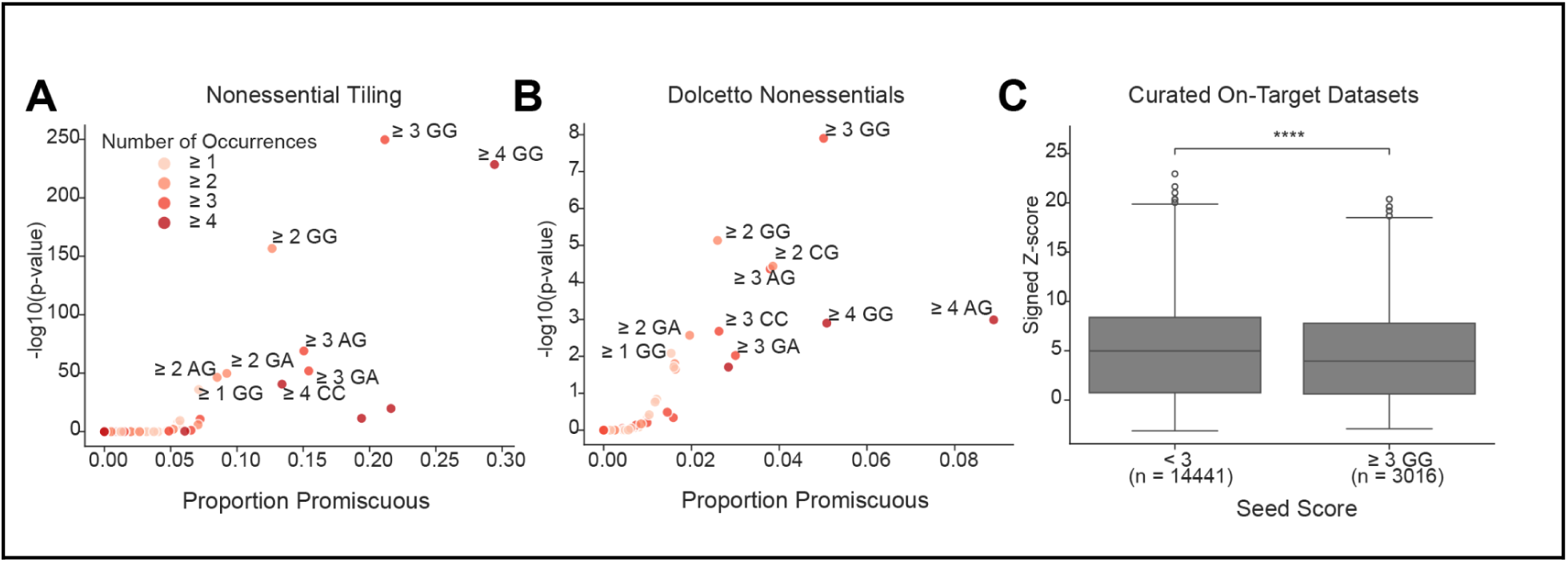
PAM-rich seed sequences promote increased CRISPRi promiscuity. (A) Scatterplot showing groups of nonessential-targeting and intergenic guides with at least a certain number of occurrences within the 12bp seed region (indicated by color) of each dinucleotide sequence. The x axis shows the proportion of guides within the group that are promiscuous, and the y axis shows p-values derived from a one-sided Fisher’s Exact test assessing the relationship between guide presence within each group (yes/no) and guide promiscuity (promiscuous/nonpromiscuous). Only groups with at least 30 guides are visualized. Points with p-value < 10^-20^ are annotated. (B) Same as (A) with nonessential-targeting guides within a Dolcetto screen. Points with p-value < 0.01 are annotated. (C) Boxplots of signed z-scores for guides in curated on-target datasets within 100bp downstream of their target TSS, stratified by Seed Score. Z-scores were averaged across guides present in all 4 tiling datasets generated in this study, as well as sign-adjusted such that a more positive score corresponds to a more active guide. X axis denotes Seed Score and number of guides in each group. Significance bar indicates the result of a one-sided Mann Whitney U test (p-value = 2.7*10^-22^). Boxes show 25th (Q1), 50th (median), and 75th (Q3) percentiles, while whiskers show Q1 - 1.5*IQR and Q3 + 1.5*IQR (where IQR is Q3 - Q1).

**Supplementary Figure 7.**
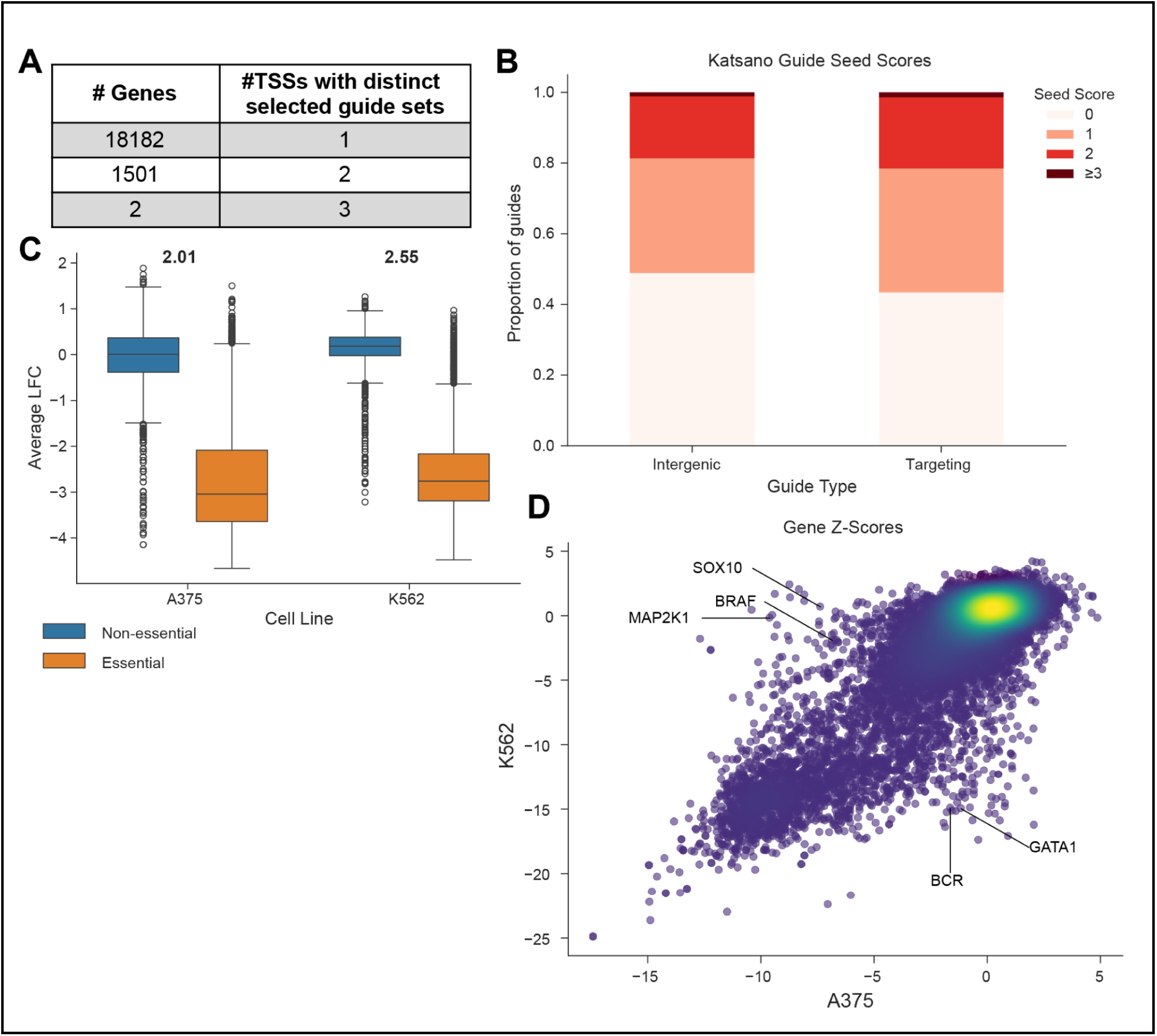
Design and evaluation of an optimized genome-wide Cas9 CRISPRi library. (A) Distribution of TSSs per gene targeted by distinct guide sets in Katsano. (B) Proportion of targeting and intergenic guides in Katsano with each Seed Score. (C) Boxplot showing average log-fold changes of guides in Katsano screens performed in A375s and K562s. Gene essentiality is indicated by color. Strictly standardized mean difference (SSMD) between non-essential and essential targeting LFCs in each screen is annotated above corresponding boxplots. Boxes show 25th (Q1), 50th (median), and 75th (Q3) percentiles, while whiskers show Q1 - 1.5*IQR and Q3 + 1.5*IQR (where IQR is Q3 - Q1). (D) Scatterplot comparing gene-level z-scores (calculated using Stouffer’s method to aggregate guide-level z-scores) in Katsano screens in A375 and K562 cell lines. Dependencies that deplete in only one cell line are labeled.

## REFERENCES

1. Doench, J.G. (2018) Am I ready for CRISPR? A user’s guide to genetic screens. Nat. Rev. Genet., 19, 67–80.

2. Przybyla, L. and Gilbert, L.A. (2022) A new era in functional genomics screens. Nat. Rev. Genet., 23, 89–103.

3. Gilbert, L.A., Larson, M.H., Morsut, L., Liu, Z., Brar, G.A., Torres, S.E., Stern-Ginossar, N., Brandman, O., Whitehead, E.H., Doudna, J.A., et al. (2013) CRISPR-mediated modular RNA-guided regulation of transcription in eukaryotes. Cell, 154, 442–451.

4. Qi, L.S., Larson, M.H., Gilbert, L.A., Doudna, J.A., Weissman, J.S., Arkin, A.P. and Lim, W.A. (2013) Repurposing CRISPR as an RNA-guided platform for sequence-specific control of gene expression. Cell, 152, 1173–1183.

5. Gilbert, L.A., Horlbeck, M.A., Adamson, B., Villalta, J.E., Chen, Y., Whitehead, E.H., Guimaraes, C., Panning, B., Ploegh, H.L., Bassik, M.C., et al. (2014) Genome-Scale CRISPR-Mediated Control of Gene Repression and Activation. Cell, 159, 647–661.

6. Horlbeck, M.A., Gilbert, L.A., Villalta, J.E., Adamson, B., Pak, R.A., Chen, Y., Fields, A.P., Park, C.Y., Corn, J.E., Kampmann, M., et al. (2016) Compact and highly active next-generation libraries for CRISPR-mediated gene repression and activation. Elife, 5.

7. Sanson, K.R., Hanna, R.E., Hegde, M., Donovan, K.F., Strand, C., Sullender, M.E., Vaimberg, E.W., Goodale, A., Root, D.E., Piccioni, F., et al. (2018) Optimized libraries for CRISPR-Cas9 genetic screens with multiple modalities. Nat. Commun., 9, 5416.

8. Jost, M., Santos, D.A., Saunders, R.A., Horlbeck, M.A., Hawkins, J.S., Scaria, S.M., Norman, T.M., Hussmann, J.A., Liem, C.R., Gross, C.A., et al. (2020) Titrating gene expression using libraries of systematically attenuated CRISPR guide RNAs. Nat. Biotechnol., 38, 355–364.

9. Dixit, A., Parnas, O., Li, B., Chen, J., Fulco, C.P., Jerby-Arnon, L., Marjanovic, N.D., Dionne, D., Burks, T., Raychowdhury, R., et al. (2016) Perturb-seq: Dissecting molecular circuits with scalable single-cell RNA profiling of pooled genetic screens. Cell, 167, 1853–1866.e17.

10. Adamson, B., Norman, T.M., Jost, M., Cho, M.Y., Nuñez, J.K., Chen, Y., Villalta, J.E., Gilbert, L.A., Horlbeck, M.A., Hein, M.Y., et al. (2016) A multiplexed single-cell CRISPR screening platform enables systematic dissection of the unfolded protein response. Cell, 167, 1867–1882.e21.

11. Datlinger, P., Rendeiro, A.F., Schmidl, C., Krausgruber, T., Traxler, P., Klughammer, J., Schuster, L.C., Kuchler, A., Alpar, D. and Bock, C. (2017) Pooled CRISPR screening with single-cell transcriptome readout. Nat. Methods, 14, 297–301.

12. Funk, L., Su, K.-C., Ly, J., Feldman, D., Singh, A., Moodie, B., Blainey, P.C. and Cheeseman, I.M. (2022) The phenotypic landscape of essential human genes. Cell, 185, 4634–4653.e22.

13. Feldman, D., Singh, A., Schmid-Burgk, J.L., Carlson, R.J., Mezger, A., Garrity, A.J., Zhang, F. and Blainey, P.C. (2019) Optical pooled screens in human cells. Cell, 179, 787–799.e17.

14. Hanna, R.E. and Doench, J.G. (2020) Design and analysis of CRISPR–Cas experiments. Nat. Biotechnol., 38, 813–823.

15. Radzisheuskaya, A., Shlyueva, D., Müller, I. and Helin, K. (2016) Optimizing sgRNA position markedly improves the efficiency of CRISPR/dCas9-mediated transcriptional repression. Nucleic Acids Res., 44, e141.

16. Horlbeck, M.A., Witkowsky, L.B., Guglielmi, B., Replogle, J.M., Gilbert, L.A., Villalta, J.E., Torigoe, S.E., Tjian, R. and Weissman, J.S. (2016) Nucleosomes impede Cas9 access to DNA in vivo and in vitro. Elife, 5, e12677.

17. FANTOM Consortium and the RIKEN PMI and CLST (DGT), Forrest, A.R.R., Kawaji, H., Rehli, M., Baillie, J.K., de Hoon, M.J.L., Haberle, V., Lassmann, T., Kulakovskiy, I.V., Lizio, M., et al. (2014) A promoter-level mammalian expression atlas. Nature, 507, 462–470.

18. Morales, J., Pujar, S., Loveland, J.E., Astashyn, A., Bennett, R., Berry, A., Cox, E., Davidson, C., Ermolaeva, O., Farrell, C.M., et al. (2022) A joint NCBI and EMBL-EBI transcript set for clinical genomics and research. Nature, 604, 310–315.

19. ENCODE Project Consortium (2012) An integrated encyclopedia of DNA elements in the human genome. Nature, 489, 57–74.

20. DeWeirdt, P.C., McGee, A.V., Zheng, F., Nwolah, I., Hegde, M. and Doench, J.G. (2022) Accounting for small variations in the tracrRNA sequence improves sgRNA activity predictions for CRISPR screening. Nat. Commun., 13, 5255.

21. Alerasool, N., Segal, D., Lee, H. and Taipale, M. (2020) An efficient KRAB domain for CRISPRi applications in human cells. Nat. Methods, 17, 1093–1096.

22. Tycko, J., DelRosso, N., Hess, G.T., Aradhana, Banerjee, A., Mukund, A., Van, M.V., Ego, B.K., Yao, D., Spees, K., et al. (2020) High-Throughput Discovery and Characterization of Human Transcriptional Effectors. Cell, 183, 2020–2035.e16.

23. Kristof, A., Karunakaran, K., Allen, C., Mizote, P., Briggs, S., Jian, Z., Nash, P. and Blazeck, J. (2025) Engineering novel CRISPRi repressors for highly efficient mammalian gene regulation. Genome Biol., 26, 164.

24. Kuscu, C., Arslan, S., Singh, R., Thorpe, J. and Adli, M. (2014) Genome-wide analysis reveals characteristics of off-target sites bound by the Cas9 endonuclease. Nat. Biotechnol., 32, 677–683.

25. Wu, X., Scott, D.A., Kriz, A.J., Chiu, A.C., Hsu, P.D., Dadon, D.B., Cheng, A.W., Trevino, A.E., Konermann, S., Chen, S., et al. (2014) Genome-wide binding of the CRISPR endonuclease Cas9 in mammalian cells. Nat. Biotechnol., 32, 670–676.

26. Rohatgi, N., Fortin, J.-P., Lau, T., Ying, Y., Zhang, Y., Lee, B.L., Costa, M.R. and Reja, R. (2024) Seed sequences mediate off-target activity in the CRISPR-interference system. Cell Genom.

27. Southard, K.M., Ardy, R.C., Tang, A., O’Sullivan, D.D., Metzner, E., Guruvayurappan, K. and Norman, T.M. (2024) Comprehensive transcription factor perturbations recapitulate fibroblast transcriptional states. bioRxivorg, 10.1101/2024.07.31.606073.

28. McGee, A.V., Liu, Y.V., Griffith, A.L., Szegletes, Z.M., Wen, B., Kraus, C., Miller, N.W., Steger, R.J., Escude Velasco, B., Bosch, J.A., et al. (2024) Modular vector assembly enables rapid assessment of emerging CRISPR technologies. Cell Genom., 4, 100519.

29. Griffith, A.L., Zheng, F., McGee, A.V., Miller, N.W., Szegletes, Z.M., Reint, G., Gademann, F., Nwolah, I., Hegde, M., Liu, Y.V., et al. (2023) Optimization of Cas12a for multiplexed genome-scale transcriptional activation. Cell Genom., 3, 100387.

30. Nuñez, J.K., Chen, J., Pommier, G.C., Cogan, J.Z., Replogle, J.M., Adriaens, C., Ramadoss, G.N., Shi, Q., Hung, K.L., Samelson, A.J., et al. (2021) Genome-wide programmable transcriptional memory by CRISPR-based epigenome editing. Cell, 184, 2503–2519.e17.

31. Hart, T., Chandrashekhar, M., Aregger, M., Steinhart, Z., Brown, K.R., MacLeod, G., Mis, M., Zimmermann, M., Fradet-Turcotte, A., Sun, S., et al. (2015) High-resolution CRISPR screens reveal fitness genes and genotype-specific cancer liabilities. Cell, 163, 1515–1526.

32. Bassik, M.C., Kampmann, M., Lebbink, R.J., Wang, S., Hein, M.Y., Poser, I., Weibezahn, J., Horlbeck, M.A., Chen, S., Mann, M., et al. (2013) A systematic mammalian genetic interaction map reveals pathways underlying ricin susceptibility. Cell, 152, 909–922.

33. Klein, D.C. and Hainer, S.J. (2020) Genomic methods in profiling DNA accessibility and factor localization. Chromosome Res., 28, 69–85.

34. Drepanos, L.M., Srikanth, S., Kaplan, E.G., Shah, S.T., Velasco, B.E., Merzouk, S. and Doench, J.G. (2025) Balancing off-target and on-target considerations for optimized Cas9 CRISPR knockout library design. bioRxiv, 10.1101/2025.08.26.672375.

35. Doench, J.G., Fusi, N., Sullender, M., Hegde, M., Vaimberg, E.W., Donovan, K.F., Smith, I., Tothova, Z., Wilen, C., Orchard, R., et al. (2016) Optimized sgRNA design to maximize activity and minimize off-target effects of CRISPR-Cas9. Nat. Biotechnol., 34, 184–191.

36. Tsai, S.Q., Zheng, Z., Nguyen, N.T., Liebers, M., Topkar, V.V., Thapar, V., Wyvekens, N., Khayter, C., Iafrate, A.J., Le, L.P., et al. (2015) GUIDE-seq enables genome-wide profiling of off-target cleavage by CRISPR-Cas nucleases. Nat. Biotechnol., 33, 187–197.

37. Rosenbluh, J., Xu, H., Harrington, W., Gill, S., Wang, X., Vazquez, F., Root, D.E., Tsherniak, A. and Hahn, W.C. (2017) Complementary information derived from CRISPR Cas9 mediated gene deletion and suppression. Nat. Commun., 8, 15403.

38. Replogle, J.M., Saunders, R.A., Pogson, A.N., Hussmann, J.A., Lenail, A., Guna, A., Mascibroda, L., Wagner, E.J., Adelman, K., Lithwick-Yanai, G., et al. (2022) Mapping information-rich genotype-phenotype landscapes with genome-scale Perturb-seq. Cell, 185, 2559–2575.e28.

39. Jaganathan, K., Ersaro, N., Novakovsky, G., Wang, Y., James, T., Schwartzentruber, J., Fiziev, P., Kassam, I., Cao, F., Hawe, J., et al. (2025) Predicting expression-altering promoter mutations with deep learning. Science.

40. Misek, S.A., Fultineer, A., Kalfon, J., Noorbakhsh, J., Boyle, I., Roy, P., Dempster, J., Petronio, L., Huang, K., Saadat, A., et al. (2024) Germline variation contributes to false negatives in CRISPR-based experiments with varying burden across ancestries. Nat. Commun., 15, 4892.

41. Dudnyk, K., Cai, D., Shi, C., Xu, J. and Zhou, J. (2024) Sequence basis of transcription initiation in the human genome. Science, 384, eadj0116.

42. Hart, T., Brown, K.R., Sircoulomb, F., Rottapel, R. and Moffat, J. (2014) Measuring error rates in genomic perturbation screens: gold standards for human functional genomics. Mol. Syst. Biol., 10, 733.

