## Supplementary material for "Optimized parameters for Cas9 CRISPR interference library design": Description of Supplementary Data Files.docx

**Title**: Supplementary Data 1

**Filename**: CombinedCleanedDatasets.csv

**Description**: Guide metadata for the combined tiling datasets used to assess predictive features of CRISPRi guide on-target efficacy and train Rule Set 3i, described in Figure 3a. sgRNA “cut” positions are defined as the position of the sgRNA sequence 3 base pairs from the “N” of the PAM.

**Title**: Supplementary Data 2

**Filename**: GW_ensembl_protein_coding_df_1kb.csv

**Description**: A list of genes that overlap within a 1000 base pair window in one of the 6 ways illustrated in Figure 6f. The “overlap type” column indicates the orientation in which the two genes overlap as labeled in Figure 6f. Overlap was calculated from genome-wide transcript annotations obtained from Ensembl Biomart on January 3^rd^, 2024.

**Title**: Supplementary Data 3

**Filename**: GW_mane_protein_coding_df_1kb.csv

**Description**: A list of genes for which MANE Select transcripts overlap in one of the 6 ways illustrated in Figure 6f. The “overlap type” column indicates the orientation in which the two genes overlap as labeled in Figure 6f. Overlap was calculated from genome-wide transcript annotations obtained from Ensembl Biomart on January 3rd, 2024, and MANE Select transcript IDs were obtained from GENCODE48 annotations.

**Title**: Supplementary Data 4

**Filename**: Katsano_Picking_Progressions.csv

**Description**: A table reporting the criteria used to prioritize and select guide RNAs for the Katsano library. gnomAD variable sites were defined as those with a variant frequency greater than 5% across the total human population or greater than 12.5% within the African American population subgroup.

**Title**: Supplementary Data 5

**Filename**: Reagents.csv

**Description**: Table of key reagents and resources used in this study.

**Title**: Supplementary Data 6

**Filename**: katsano_4_guide_per_tss_designs.csv

**Description**: Guides selected with the criteria used to design the Katsano library, including four guides per target transcription start site (TSS). The “Selection Category” column indicates why the TSS for which each particular guide was selected was chosen as a target, with “Ensembl Canonical” indicating that it belongs to an Ensembl Canonical transcript and other categories indicating that the transcript was nominated by the Jaganathan et al. transcript set. The “Pick Order” column indicates the order in which the guides were chosen.

**Title**: Supplementary Data 7

**Filename**: katsano_guide_gene_mapping_GENCODE48.csv

**Description**: Guide annotations for the Katsano library with three guides per target. Every TSS in the original Katsano target set that each guide in the library targets is included as a separate row. A guide was considered to target a TSS if it falls within [-50,300] base pairs of it.
